# A patient-centric therapeutic paradigm uncouples prostate cancer suppression from systemic metabolic collapse

**DOI:** 10.64898/2026.08.30.748180

**Authors:** Ziqun Liu, Hanchu Ye, Haoran Hu, Wuhong Chen, Chen Wang, Yongfeng Yang, Xinran Yan, Nathan longlong Chen, Yang Wang, Yumeng Yang, Chunqiao Sun, Yaqing Bai, Haiyang Hou, Xinan Wang, Jun Cao, Qinsheng Chen, Qi Wang, Yuting Xiao, Yuanyuan Cai, Jiahui Chen, Feifei Lin, Yushi Li, Yuanhao Dang, Yuxin Su, Xiaofei Wang, Zheng Tian, Yidong Shen, Linfeng Sun, Jia Liu, Denglong Wu, Chengchao Ruan, Ruobing Ren, Youhong Hu, Huiru Tang, Shengsong Huang, Zhenfei Li

## Abstract

The clinical benefits of cancer therapies are often compromised by the ‘tolerable’ adverse effects that impair systemic organismal health and may evolve into latent life threats. Here, we identified profound abiraterone-induced but androgen-independent metabolic perturbations in prostate cancer patients and developed Lifehug-9892 to balance tumor therapy with systemic metabolic homeostasis. By integrating population cohorts with high-resolution metabolomics, we demonstrate that abiraterone induces profound systemic lipidomic dysregulation, characterized by the massive, pathological accumulation of desmosterol. Abiraterone inhibits but stabilizes DHCR24, leading to a metabolic trap in patients showing elevated levels of both desmosterol and cholesterol. Desmosterol accumulation is highly lipotoxic, potently triggering endothelial cell senescence and necrosis, macrophage foam cell formation, murine atherosclerosis, and hepatic senescence. To mechanistically uncouple and therapeutically rescue this systemic metabolic collapse, Lifehug-9892 was rationally designed to selectively retain on-target CYP17A1 inhibition while completely sparing DHCR24 function. Lifehug-9892 maintains potent tumor-suppressive activity while fully preserving the desmosterol-cholesterol metabolic axis and preventing systemic cardiovascular and hepatic damage. Our study uncovers a critical mechanistic link between drug-induced metabolic dysregulation and organismal health in cancer patients, providing a biochemical framework for developing patient-centric targeted therapies that preserve host homeostasis.

**One Sentence Summary:** Abiraterone inhibits DHCR24 to induce systemic metabolic dysregulation in prostate cancer patients.

## Introduction

A critical question in mordent oncology is understanding the drivers of mortality when patients do not succumb directly to their cancer. The primary focus of cancer therapy has long been eradicating cancer cells, with relatively less emphasis on the long-term systemic toxicity of drugs (1, 2). Side effects, though marked as ‘tolerable’, are indicators of impaired organismal health and might provide hints for patient mortality. Elucidating the molecular mechanisms underlying drug-induced organismal damage would guide drug optimization and disease management.

The clinical trajectory of next-generation androgen receptor pathway inhibitors (ARPIs), such as abiraterone and enzalutamide, perfectly illustrates this paradigm. Developed to treat castration-resistant prostate cancer, ARPIs are increasingly prescribed for earlier-stage disease due to their potent tumor-suppressive capabilities (3–5). However, a stark clinical paradox has emerged: ARPIs exhibit superior efficacy in inhibiting disease progression than in actually prolonging overall survival—a discrepancy that becomes profoundly magnified when these drugs are administered in earlier disease stages (NCT00638693, NCT00887198, and NCT00269476 for abiraterone; NCT00974311 and NCT01212991 for enzalutamide) (6–11). We hypothesize that this paradox is driven by drug-induced systemic metabolic impairment. Crucially, the severe adverse events associated with ARPIs—specifically the progressive cardiovascular and hepatic disorders predominantly observed with abiraterone (6–9) —are highly drug-specific. Because these distinct pathophysiologies do not overlap across the ARPI class, they are not inevitable consequences of suppressing the androgen axis. Instead, this clinical divergence strongly implicates distinct, off-target pharmacological interactions, providing a definitive mechanistic rationale for circumventing these toxicities via rational drug redesign.

To define this metabolic vulnerability, we investigated abiraterone-induced systemic metabolomic alterations in prostate cancer patients. Here, we reveal that abiraterone acts as an unintended inhibitor of DHCR24, driving profound, androgen-independent lipidomic perturbations. Guided by these mechanistic insights, we developed Lifehug-9892, a rationally engineered compound that meticulously uncouples anti-tumor efficacy from systemic toxicity, effectively suppressing tumor growth while fully preserving the host’s metabolic homeostasis.

## Results

### Abiraterone induces systemic metabolic dysregulation

To explore potential abiraterone-induced systemic perturbations in patients, serum samples were retrospectively collected from a cohort of 15 metastatic hormone-sensitive prostate cancer (mHSPC) patients receiving androgen deprivation therapy (ADT) combined with abiraterone-prednisone treatment. Samples obtained at baseline as well as at 1, 2, 3, and more than 7 months post-treatment were subjected to comprehensive metabolomics analyses, including targeted lipidomics, derivatized sterol profiling, and untargeted metabolomics. A total of 1375 lipid molecules, 31 sterols, and 1470 polar metabolites were detected with these clinical specimens (**Fig. 1A**). Abiraterone effectively reduced serum concentrations of dehydroepiandrosterone (DHEA) and cortisol, as anticipated (**Fig. S1A**). Surprisingly, desmosterol exhibited the most profound increase, and its accumulation was observed at all post-treatment time points and in every patient (**Fig. 1 A and B**). Serum cholesterol levels exhibited a mildly increasing trend, and showed a statistically significant upregulation after 3 months of abiraterone treatment (**Fig. 1C**). Alterations in lipidomics profiles were also observed. Most of the serum lipids showed steady upward trends after abiraterone treatment (**Fig. S1B**). Subclass analysis revealed that phosphatidylcholine (PC) was the most significant enriched lipid subclass after abiraterone treatment. Phosphatidylethanolamine (PE), phosphatidylserine (PS), and other glycerophospholipids were also significantly enriched **(Fig. S1C and D).** Together, these results demonstrate systemic perturbations in abiraterone-treated patients.

**Fig. 1.**
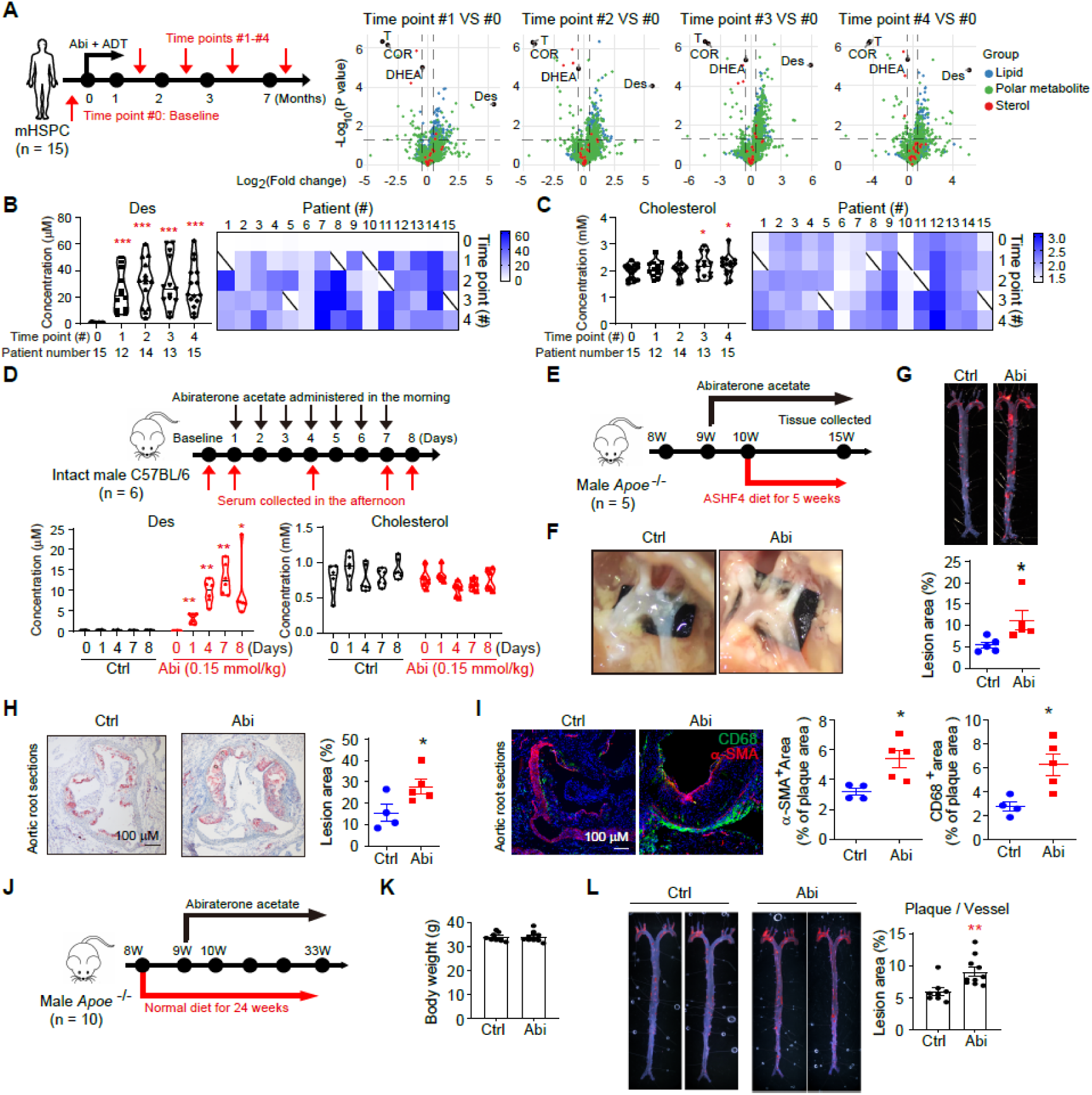
Abiraterone induces systemic metabolic perturbations in patients and mice. A, Schematic overview of clinical specimens collection and serum metabolomic profiling of prostate cancer patients receiving abiraterone (Abi) treatment. Metastatic hormone-sensitive prostate cancer (mHSPC) patients (n = 15) initiate androgen deprivation therapy (ADT) and abiraterone acetate/prednisone treatment at time point #0. Des, desmosterol; DHEA, dehydroepiandrosterone; T, testosterone; COR, cortisol. **B and C**, Alterations of serum desmosterol (B) and cholesterol (C) concentration in patients. Heatmap showing the dynamic alterations for each individual patient. **D**, Dynamic alterations of desmosterol and cholesterol in abiraterone acetate (0.15 mmol/kg/day; Intraperitoneal injection) treated mice (n = 6 per group). Intact male C57BL/6 mice were treated with abiraterone acetate alone. **E**, A schematic showing long term effect of abiraterone in male *Apoe*^-/-^ C57BL/6 mice fed with ASHF4 diet. Intact male mice were fed with ASHF4 diet (containing 40 kcal% fat, 1.25% cholesterol, 0.5% sodium cholate) for 5 weeks. **F**, Representative images of atherosclerotic plaques in the aorta arch from ASHF4 diet fed male *Apoe*^-/-^ mice treated with or without abiraterone acetate. **G**, Oil Red O staining of en face aorta from ASHF4 diet fed male *Apoe*^-/-^ mice treated with or without abiraterone acetate (n = 5 per group). **H**, Oil Red O staining of aortic root sections from ASHF4 diet fed male *Apoe*^-/-^ mice treated with or without abiraterone acetate (Ctrl group, n = 4; Abi group, n = 5). **I**, Immunofluorescence staining of α-SMA (red), CD68 (green), and DAPI (blue) in aortic root sections from ASHF4 diet fed male *Apoe* ^-/-^ mice (Ctrl group, n = 4; Abi group, n = 5). **J**, A schematic showing abiraterone function in male *Apoe* ^-/-^ mice fed with normal diet (Ctrl group, n = 8; Abi group, n = 10). **K**, Body weight of normal diet fed male *Apoe* ^-/-^ mice treated with or without abiraterone acetate (Ctrl group, n = 8; Abi group, n = 10). **L**, Atherosclerotic plaques of en face aorta in normal diet fed male *Apoe*^-/-^ mice (Ctrl group, n = 8; Abi group, n = 10). Data are shown as the mean ± SEM. *, ** and *** denoted P < 0.05, P < 0.01 and P < 0.001, respectively.

ADT has been reported to perturb cholesterol synthesis and lipid metabolism (12, 13). To exclude the potential interference of ADT on systemic metabolic alterations, intact male mice were treated with abiraterone alone (0.15 mmol/kg/day, ≈ 58.65 mg/kg/day; without prednisone) for 7 consecutive days, and serum samples were collected at the baseline time point as well as 1, 4, 7, and 8 days after treatment initiation (**Fig. 1D**). Since mice lack adrenal CYP17A1 expression, abiraterone has a limited impact on murine steroidogenesis (**Fig. S2A**). Without disturbing steroidogenesis, abiraterone induced desmosterol accumulation immediately following the first dose. Desmosterol concentration increased continuously with abiraterone administration, but dropped following abiraterone withdrawal (**Fig. 1D**). Cholesterol concentration was not altered throughout the abiraterone treatment period. Additionally, abiraterone treatment led to alterations in lipid profiles (**Fig. S2 B and C**). Collectively, these results demonstrate that abiraterone directly induces systemic metabolic dysregulation.

### Abiraterone accelerates cardiovascular disease

The pathological effects of the abiraterone-induced systemic metabolism alterations were further investigated. Considering the well-established correlation between cholesterol-lipid metabolism and cardiovascular disease, *Apoe*^-/-^ mice (Apolipoprotein E knockout mice, a model for atherosclerosis) fed with a high-cholesterol, high-fat (ASHF4) diet were treated with abiraterone for 6 weeks (**Fig. 1E**). An increase of atherosclerotic plaque in the aortic arch (**Fig. 1F**) and whole aorta (**Fig. 1G**) was observed in abiraterone-treated mice. Moreover, aortic root analysis of abiraterone-treated *Apoe^-/-^* mice showed increased lipid deposition (**Fig. 1H**). Immunofluorescent analyses demonstrated that plaques in the abiraterone-treated *Apoe*^-/-^ mice contained increased myofibroblast activation and macrophage infiltration (**Fig. 1I**). To more accurately simulate clinical conditions in patients, a normal diet was given to *Apoe*^-/-^mice (**Fig. 1J**). Similarly, abiraterone accelerated plaque generation, although a longer treatment duration was required **(Fig. 1 K and L**). To determine whether the cholesterol metabolism pathway is involved in this pathological effect of abiraterone, rosuvastatin calcium (5 mg/kg/day, per os) or ezetimibe (10 mg/kg/day, per os) was administered to *Apoe*^-/-^ mice to block cholesterol synthesis or absorption, respectively(14) (15). Rosuvastatin blocked abiraterone-induced plaque generation, whereas ezetimibe showed a more potent inhibition (**Fig. S3**). These results together demonstrate that abiraterone promotes the progression of cardiovascular disease.

### Abiraterone acts as an off-target inhibitor and paradoxical stabilizer of DHCR24

To explore the mechanisms underlying abiraterone-mediated dysregulation of cholesterol metabolism, virtual screening was performed using Boltzmann 2 (Boltz-2) to determine the affinity of abiraterone for all sterol metabolism-related enzymes (16). DHCR24 emerged as one of the top candidates, with a direct link to cholesterol metabolism (**Fig. 2A**). Additionally, the immediate and dramatic accumulation of desmosterol in abiraterone-treated mice also suggested a direct biochemical inhibition of DHCR24 by abiraterone. To confirm this direct inhibition, an in vitro DHCR24 activity assay was conducted in the presence or absence of abiraterone. Abiraterone suppressed cholesterol production when DHCR24 was incubated with desmosterol as the substrate (**Fig. 2B**). Abiraterone also inhibited DHCR24 activity in VCaP, Huh7, and PC3 cells, as indicated by the accumulated intracellular desmosterol after abiraterone treatment (**Fig. 2C**). Direct binding of abiraterone and its downstream metabolites to purified DHCR24 protein was verified by surface plasmon resonance (SPR). Abiraterone exhibited strong affinity to DHCR24, whereas the 5α-reduced downstream metabolites (3α-OH-5α-Abi and 3β-OH-5α-Abi) exhibited higher affinity for DHCR24 than the 5β-reduced metabolites (3α-OH-5β-Abi and 3β-OH-5β-Abi) (**Fig. 2D; Fig. S4A**)(17, 18). Consistently, administration of delta-4-abiraterone (D4A) and 5α-reduced abiraterone (5α-Abi) to mice showed no consistent alterations of testosterone and corticosterone, but markedly induced desmosterol accumulation comparable to that induced by abiraterone, while 5β-reduced abiraterone (5β-Abi) had almost no effect on desmosterol accumulation (**Fig. 2E; Fig. S4B**). These results together provide the mechanistic insights into desmosterol accumulation following abiraterone treatment in both patient cohorts and mouse models.

**Fig. 2.**
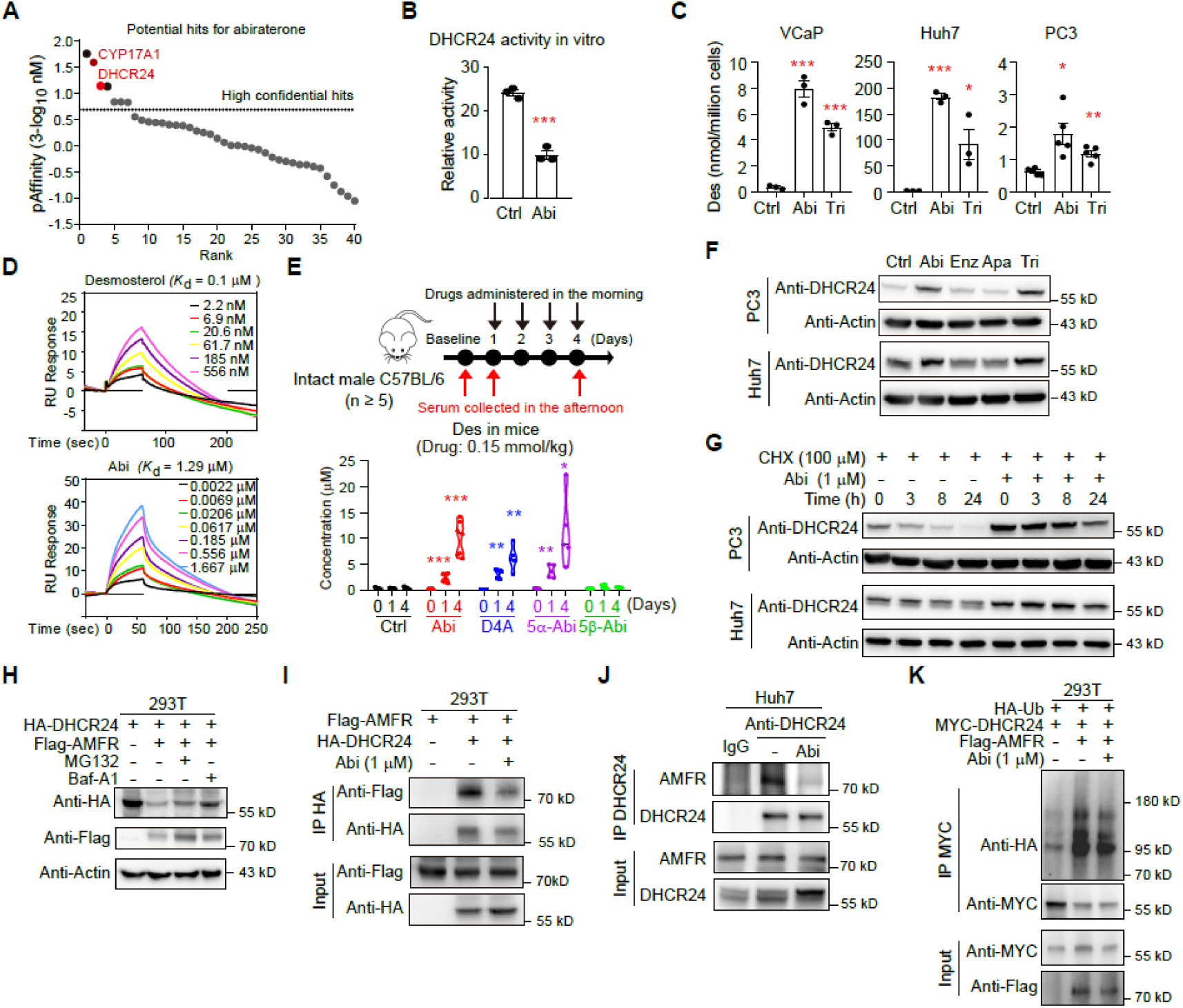
Abiraterone acts as an off-target DHCR24 inhibitor. **A**, Potential hits for abiraterone predicted by Boltz-2 across sterol-metabolism enzymes. Each dot represents one target protein; the y-axis shows pAffinity defined as 3-log_10_(Affinity [nM]), where Affinity is the Boltz-2-predicted binding affinity in nM. Targets are ranked on the x-axis from the strongest to the weakest predicted binding. The dashed line indicates the 1 μM threshold (pAffinity = 0). CYP17A1 and DHCR24 are highlighted in red as top-ranking candidates. **B**, Abiraterone inhibits DHCR24 activity in vitro. **C**, Abiraterone inhibits DHCR24 activity in different cell lines. Des, desmosterol; Tri, triparanol, a reported DHCR24 inhibitor. **D**, Surface plasmon resonance (SPR) analysis on the affinity of desmosterol and abiraterone to purified DHCR24 proteins. **E**, Desmosterol accumulation in mice treated with abiraterone and its metabolites. Intact male C57BL/6 mice were treated with abiraterone acetate or its metabolites (0.15 mmol/kg/day; Intraperitoneal injection). Serum was collected after one or four days post treatment (n = 5 per group except for Abi group with 7 mice). D4A, delta-4 abiraterone; 5α-Abi, 5α-reduced abiraterone; 5β-Abi, 5β-reduced abiraterone. Median, 25^th^ percentile, and 75^th^ percentile are shown in violin plot; the outer shape depicts the kernel density estimation of the data distribution. **F**, Abiraterone increases the protein levels of DHCR24 in PC3 and Huh7 cells. Compounds of 1 µM were used to treat different cell lines for 48 hrs. Enz, enzalutamide; Apa, apalutamide. **G**, Abiraterone prevents endogenous DHCR24 degradation. Abiraterone, 1 μM; cycloheximide (CHX), 100 μM. **H**, AMFR degrades DHCR24. MG132, proteasome inhibitor, 10 μM; bafilomycin A1 (Baf-A1), lysosome inhibitor, 100 nM. **I**, Abiraterone prevents the binding of overexpressed-DHCR24 to AMFR in 293T cells. Abiraterone, 1 μM. **J**, Abiraterone prevents the endogenous binding of DHCR24 to AMFR in Huh7 cells. Abiraterone, 1 μM. **K**, Abiraterone suppresses AMFR-induced DHCR24 ubiquitination in 293T cells. Data are shown as the mean ± SEM. *, ** and *** denoted P < 0.05, P < 0.01 and P < 0.001, respectively.

However, the eventual accumulation of cholesterol observed in patients contradicts a simple, continuous DHCR24 blockade. To explain this paradox, mechanisms for abiraterone regulating DHCR24 were further investigated. Abiraterone increased DHCR24 protein abundance in Huh7 and PC3 cells (**Fig. 2F**). Notably, abiraterone had no effect on *DHCR24* mRNA levels (**Fig. S4C**), but prolonged the half-life of DHCR24 protein (**Fig. 2G**). AMFR, previously reported as a DHCR24 ubiquitin ligase, was confirmed to degrade DHCR24 in our experimental system (**Fig. 2H**). The interaction between DHCR24 and AMFR was suppressed by abiraterone both in exogenous and endogenous context (**Fig. 2 I and J**). Therefore, abiraterone prevented AMFR-mediated ubiquitination of DHCR24 (**Fig. 2K**). Collectively, abiraterone creates a paradoxical metabolic trap: abiraterone inhibits DHCR24’s enzymatic activity while simultaneously increasing its protein stability. Clinically, abiraterone is administered in the morning to align with the circadian rhythm of androgen synthesis. Its serum concentration declines at night, a time when cholesterol synthesis becomes active. Additionally, our previous studies reported accelerated clearance of abiraterone in patients following months of treatment (1). These factors together explain the gradual increase of cholesterol in abiraterone-treated patients.

### Pathological desmosterol accumulation triggers cardiovascular and hepatic dysregulation

To investigate the pathological consequences of abiraterone-induced desmosterol accumulation, HUVECs (human umbilical vein endothelial cells) were treated with different doses of desmosterol. While cholesterol showed minor effect on cell proliferation, desmosterol at concentrations of 10 µM or higher inhibited HUVEC growth (**Fig. 3A**). High concentration of desmosterol induced HUVEC necrosis (**Fig. 3 B and C**). Transcriptomic analysis revealed that desmosterol of 20 or 50 µM, but not cholesterol, activated inflammatory signaling pathways in HUVECs (**Fig. 3D**). Notably, desmosterol also activated a senescence-associated transcriptional signature in HUVECs (**Fig. 3D**)(19). Desmosterol treatment increased the expression of *CDKN1A* (a well-established senescence marker) and *IL6* (an inflammatory marker) in HUVECs (**Fig. 3E**). Additionally, desmosterol enhanced p21 expression and senescence-associated β-galactosidase (SA-β-gal) activity in a dose-dependent manner (**Fig. 3 F and G**). Conditioned medium from desmosterol- or cholesterol-treated HUVECs was further collected for proteomic analysis. Senescence related proteins were enriched following desmosterol but not cholesterol treatment (**Fig. S5 A and B**). THP-1 monocytes were further co-incubated with conditioned medium collected from desmosterol-or cholesterol-treated HUVECs. Conditioned medium from desmosterol-treated HUVECs promoted lipid droplet accumulation in THP-1 cells, thereby facilitating foam cell formation (**Fig. 3H**). Transcriptomic analysis showed that the desmosterol-treated HUVEC conditioned medium enhanced the expression of atherosclerosis-promoting genes while suppressing genes that protect against atherosclerosis in THP-1 cells (**Fig. 3I**). Consistently, the desmosterol-treated HUVEC conditioned medium increased the expression of CD36 (a protein that facilitates fatty acid uptake) and ABCA1 (a protein that facilitates cholesterol/lipids efflux) in THP-1 cells (**Fig. 3J**). These results together demonstrate that abiraterone-induced desmosterol accumulation promotes endothelial cell necrosis and senescence along with foam cell formation, thereby contributing to the pathogenesis of cardiovascular disease.

**Fig. 3.**
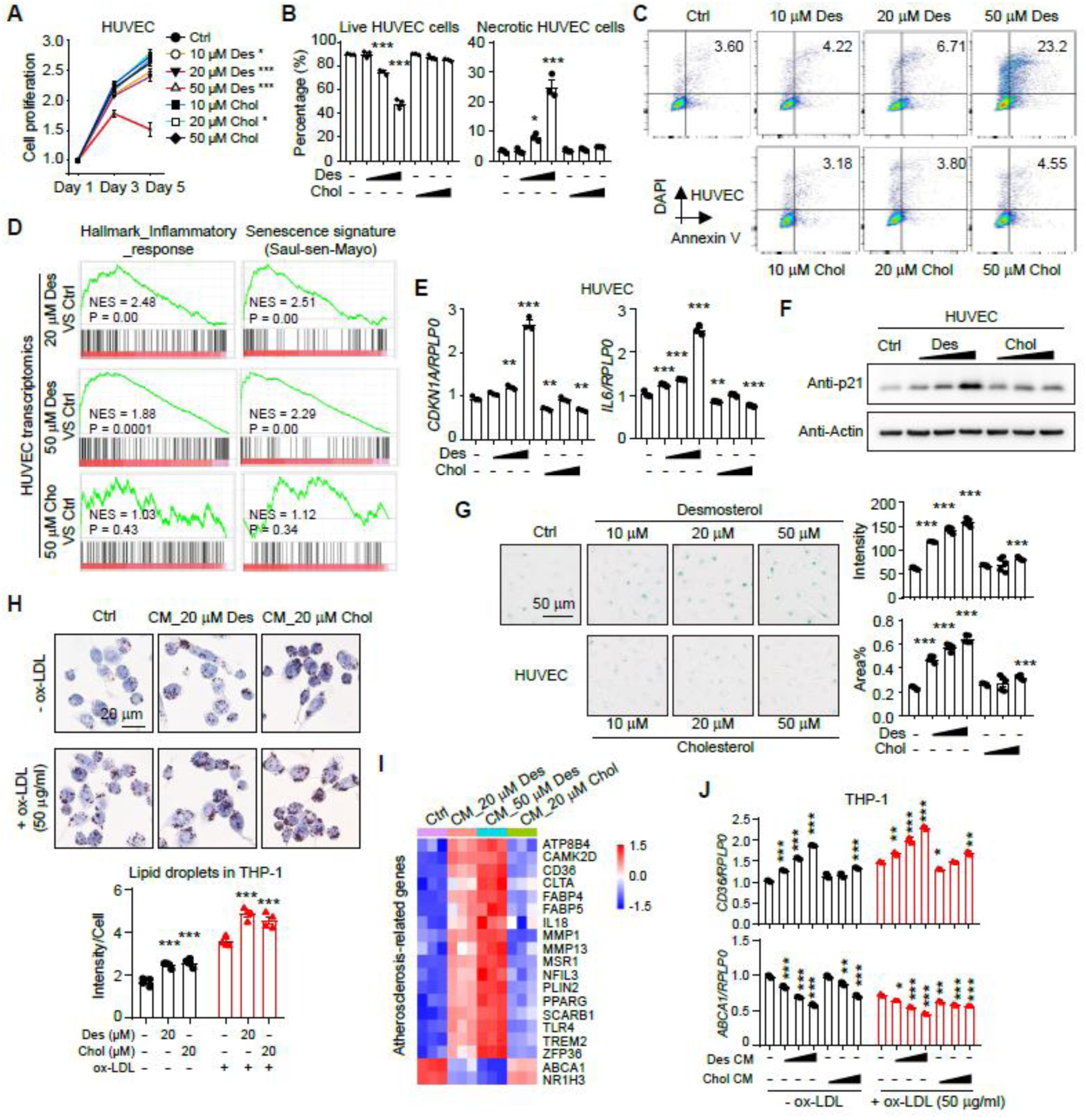
Pathological desmosterol accumulation accelerates cardiovascular disease. **A**, Viability of HUVECs treated with different doses of desmosterol and cholesterol. Des, desmosterol; Chol, cholesterol. **B and C**, High doses of desmosterol result in necrosis in HUVECs. HUVECs were treated with desmosterol (10, 20, 50 μM) or cholesterol (10, 20, 50 μM) for 72 hrs. DAPI⁺/Annexin V⁺ cells detected by flow cytometry (upper right quadrant) were classified as necrosis or late apoptosis. **D**, GSEA analysis on transcriptomics of HUVECs treated with desmosterol or cholesterol. **E**, Expression of *CDKN1A* and *IL6* in HUVECs treated with desmosterol (10, 20, 50 μM) and cholesterol (10, 20, 50 μM). **F**, Expression of p21 in HUVECs treated by desmosterol (10, 20, 50 μM) and cholesterol (10, 20, 50 μM). **G**, SA-β-gal staining in HUVECs treated by desmosterol or cholesterol. **H**, Lipid droplets in THP-1 cells treated with conditioned medium (CM) from HUVECs. CM_20 μM Des, conditioned medium from HUVECs treated by 20 μM desmosterol for 72 hrs; CM_20 μM Chol, conditioned medium from HUVECs treated by 20 μM cholesterol for 72 hrs. ox-LDL, oxidized LDL, 50 μg/ml. **I**, Expression of classical atherosclerosis-related genes in THP-1 cells after treatment with different conditioned medium for 12 hrs. **J**, Expression of *CD36* (lipid uptake receptor) and *ABCA1* (lipid efflux transporter) in THP-1 cells treated with different conditioned medium. Desmosterol or cholesterol of 10, 20, or 50 μM were used to incubate with HUVECs for 72 hrs to generate related conditioned medium. *, ** and *** denoted P < 0.05, P < 0.01 and P < 0.001, respectively.

To rule out the role of residual desmosterol in the desmosterol-treated HUVEC conditioned medium on THP-1 cell responses, the conditioned medium was extracted using methyl tert-butyl ether (MTBE). Extracts from the aqueous phase, but not the organic phase, activated *CD36* expression in THP-1 cells (**Fig. S5C**). Consistently, heat inactivation or proteinase K treatment of HUVEC conditioned medium abrogated its ability to regulate *CD36* expression in THP-1 cells. Proteins with a molecular weight exceeding 10 kDa were identified as the major functional components responsible for regulating *CD36* expression in THP-1 cells (**Fig. S5C**). These results together indicate that the senescence-associated secretory phenotype (SASP)-related proteins secreted by HUVECs but not the residual desmosterol in the conditioned medium promote foam cell formation.

The senescence phenotype observed in desmosterol-treated HUVECs attracted our attention. Huh7 and NIH/3T3 cells were also treated with desmosterol, and enhanced SA-β-gal activity and p21 expression were detected in both cell lines (**Fig. 4 A-C**). Furthermore, abiraterone also upregulated SA-β-gal activity and p21 expression in Huh7 and NIH/3T3 cells (**Fig. 4 D and E**). Since a high serum concentration of desmosterol could not be achieved via intraperitoneal injection, intravenous injection, or oral administration, we treated C57BL/6 mice with abiraterone acetate to induce desmosterol accumulation for over 18 weeks. Different organs were collected for SA-β-gal staining. The liver exhibited a markedly enhanced SA-β-gal signal following abiraterone treatment (**Fig. 4F**). Transcriptomic analysis revealed that abiraterone induced senescence and inflammatory signaling pathways in the liver (**Fig. 4G**). The upregulation of genes related to senescence and inflammation was further validated by quantitative PCR (**Fig. 4H**). To explore the correlation between abiraterone and senescence in patients, a senescence-associated signature composed of lipid metabolites was first established. This signature was established by identifying lipid metabolites that are consistently increased with ageing in a longitudinal mouse study and subsequently validated in a human ageing cohort from the Fudan University Human Phenome Project (n = 1,003; with participants aged 20 to 60 years old) (**Fig. 4 I and J**). Notably, the lipidomics-ageing signature was enriched in mice treated with abiraterone acetate (mouse lipidomics results from Fig. 1D were used for analysis) (**Fig. 4K**), and similarly enriched in serum samples from patients receiving abiraterone (patient lipidomics results from Fig. 1A were used for analysis) (**Fig. 4L**). Collectively, these data suggest that abiraterone may accelerate organismal senescence, with a particularly prominent effect in the liver.

**Fig. 4.**
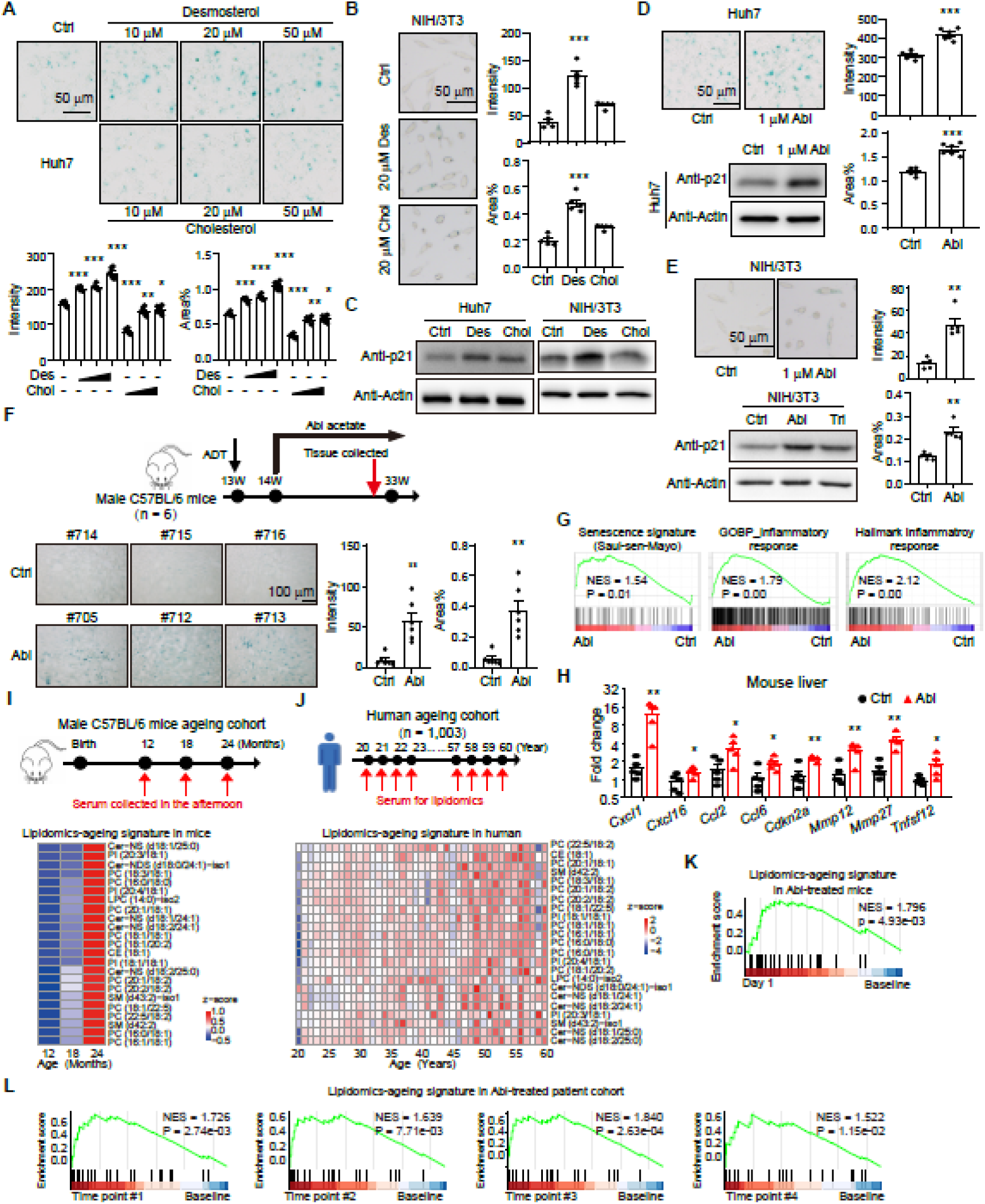
Pathological desmosterol accumulation accelerates hepatic senescence. **A**, SA-β-gal staining in Huh7 cells treated with different dose of desmosterol (Des) or cholesterol (Chol) for 24 hrs. Representative images are shown (n = 6 images per group). **B**, SA-β-gal staining in NIH/3T3 cells treated with desmosterol (20 µM) or cholesterol (20 µM) for 96 hrs. Representative images are shown (n = 5 images per group). **C**, p21 expression in Huh7 and NIH/3T3 cells. Cells were treated with 20 μM desmosterol or cholesterol for 48 hrs. **D and E**, SA-β-gal staining and p21 expression in Huh7 and NIH/3T3 cells treated with or without 1 μM abiraterone (Abi). **F**, SA-β-gal staining with liver tissues from male C57BL/6 mice treated with or without abiraterone acetate. Representative images are shown (n = 6 per group). **G**, GSEA on transcriptomics of mouse liver tissues treated with or without abiraterone (n = 5 per group). **H**, Quantitative PCR results of senescence markers in liver tissues from mice treated with or without abiraterone (n = 5 per group). **I**, Lipidomics-ageing signature in mouse cohort (12-month-old mice, n = 8; 18-month-old mice, n = 6; 24-month-old mice, n = 5). **J**, Lipidomics-ageing signature in population cohort of the Fudan University Human Phenome Project (n = 1003). **K and L**, Enrichment of lipidomics-ageing signature in abiraterone acetate treated mice (K) and patients (L). *, ** and *** denoted P < 0.05, P < 0.01 and P < 0.001, respectively.

### Rational design of LH9892

Rational drug design strategies for optimizing the abiraterone structure to efficiently inhibit CYP17A1 while evading recognition by DHCR24 would facilitate the dual outcomes of suppressing cancer cell development and maintaining organismal metabolic homeostasis. A series of abiraterone analogs were then designed and synthesized, the final compound LifeHug-9892 (LH9892) was selected after structure-activity assays. Both Boltz-2-based virtual screening and cell-based functional assays showed that LH9892 exhibited inhibitory activity comparable to that of abiraterone, while showing no effect on DHCR24 activity (**Fig. 5 A and B**). Consistently, LH9892 effectively inhibited pregnenolone-induced AR target genes expression and cell growth in VCaP cells stably expressing CYP17A1 (VCaP-CYP17A1 cells) (**Fig. 5 C and D**). The antitumor activity of LH9892 was then evaluated in mice with xenografts from VCaP-CYP17A1 cells. LH9892 efficiently suppressed pregnenolone-induced xenograft growth (**Fig. 5E**). These results together demonstrate that LH9892 retains antitumor activity equivalent to that of abiraterone.

**Fig. 5.**
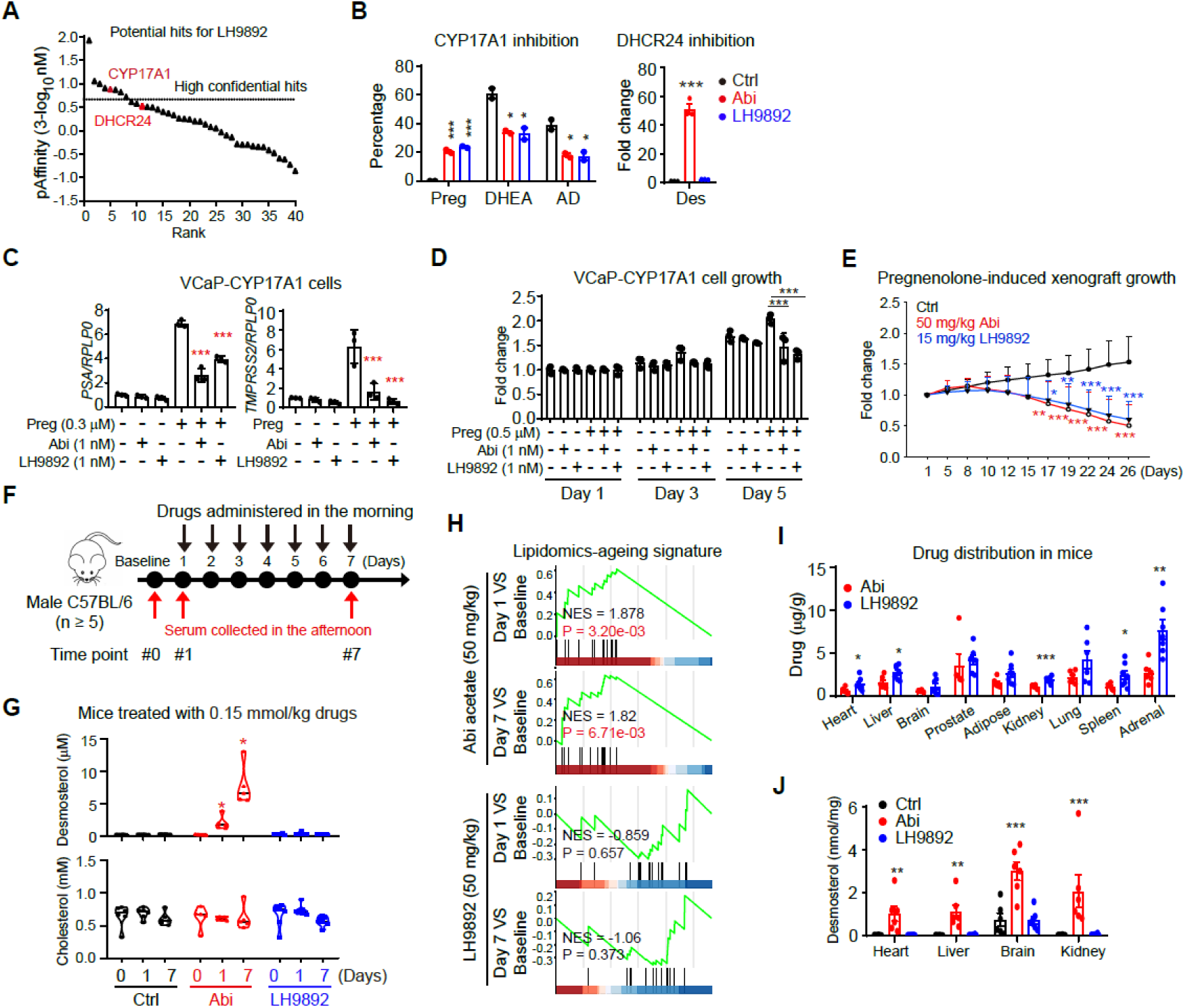
Development of Lifehug-9892 (LH9892). **A**, Potential hits for LH9892 predicted by Boltz-2 across sterol-metabolism enzymes. Targets are ranked on the x-axis from the strongest to the weakest predicted binding. The dashed line indicates the 1 μM threshold (pAffinity = 0). **B**, Effect of LH9892 on the activities of CYP17A1 and DHCR24. VCaP cells stably expressing CYP17A1 (VCaP-CYP17A1) were incubated with [^3^H]-pregnenolone with different compounds of 0.1 nM to evaluate CYP17A1 activity; intracellular desmosterol (Des) was detected by LC-MS after Huh7 cells incubated with different compounds of 1 μM for 5 hrs to evaluate DHCR24 activity. **C**, LH9892 suppresses pregnenolone-induced AR target gene expression in VCaP-CYP17A1 cells. Preg, pregnenolone, 0.3 µM. **D**, LH9892 inhibits pregnenolone-induced cell proliferation in VCaP-CYP17A1 cells. **E**, LH9892 suppresses pregnenolone-induced xenograft growth. Male NDG mice were castrated and implanted with sustained-release pregnenolone pellets. VCaP-CYP17A1 cells were implanted for xenograft assay. Abiraterone acetate, 50 mg/kg/day, per os; LH9892, 15 mg/kg/day, per os. **F**, A schematic illustrating abiraterone acetate (0.15 mmol/kg/day, intraperitoneal injection) or LH9892 (0.15 mmol/kg/day, intraperitoneal injection) treatment in mice. **G**, Dynamic alterations of desmosterol and cholesterol in abiraterone acetate or LH9892-treated mice. Violin plots showing median, 25^th^ percentile, and 75^th^ percentile with the outer shape showing the kernel density estimation of the data distribution. **H**, Enrichment of lipidomics-ageing signature in abiraterone acetate but not LH9892-treated mice. **I and J**, Drug distributions (I) and desmosterol levels (J) in different mice tissues. Tissues were collected from mice after abiraterone acetate or LH9892 treatment for 7 days. *, ** and *** denoted P < 0.05, P < 0.01 and P < 0.001, respectively.

The direct effect of LH9892 on systemic metabolic homeostasis was also evaluated. Mice were treated with abiraterone acetate (0.15 mmol/kg/day, intraperitoneal injection) or LH9892 (0.15 mmol/kg/day, intraperitoneal injection), and serum was collected for metabolomics analysis **(Fig. 5F)**. Overall, fewer metabolites were altered by LH9892, compared to abiraterone acetate (**Fig. S6 A and B**). The perturbations of systemic metabolic homeostasis were mild in LH9892-treated mice (**Fig. S6C**). For desmosterol levels, abiraterone acetate treatment induced an immediate and robust elevation in desmosterol levels, whereas LH9892 exerted no effect on desmosterol levels, even after 7 consecutive days of treatment **(Fig. 5G)**. The lipidomics-ageing signature was enriched in abiraterone-treated mice, but not in LH9892-treated mice (**Fig. 5H**). Tissues from these mice were also collected for drug distribution analysis. The tissue distribution profile of LH9892 in mice was similar to that of abiraterone, but with a higher tissue concentration (**Fig. 5I**). However, desmosterol levels in various tissues from LH9892-treated mice remained unaffected (**Fig. 5J**). These results together demonstrate that LH9892 preserves organismal metabolic homeostasis.

The pathological effects of LH9892 were further evaluated. LH9892 showed limited effect on SA-β-gal activity and p21 expression in Huh7 cells (**Fig. 6 A and B**). Furthermore, mice were treated with either abiraterone (50 mg/kg/day) or LH9892 (15 and 50 mg/kg/day) for 8 consecutive weeks (**Fig. 6C**). After prolonged treatment, neither of these two drugs exerted an obvious impact on body weight or liver weight (**Fig. 6C**). Long term treatment of abiraterone enhanced SA-β-gal activity in liver tissue. LH9892 of 15 mg/kg/day showed no effect on SA-β-gal activity, while LH9892 of 50 mg/kg/day surprisingly suppressed SA-β-gal activity (**Fig. 6D**). Liver tissues were also collected for proteomic analyses. A proteomic senescence signature (Saul-sen-Mayo senescence signature) was enriched in liver tissues from abiraterone-treated mice, but not in those from LH9892-treated mice (**Fig. 6E**)(19). Consistently, the lipidomics-ageing signature was enriched in serum lipidomics from abiraterone-treated mice, but not those from LH9892-treated mice (**Fig. 6F**). The effect of LH9892 on atherosclerosis was also evaluated. *Apoe*^-/-^mice fed with ASHF4 diet were treated with LH9892 for 9 weeks (**Fig. 6G**). The results showed that LH9892 promoted the increase of body weight, but decreased the atherosclerotic lesions in aortic arch and whole aorta (**Fig. 6H**). Together, these data confirm that LH9892 avoids the adverse effect on hepatic and cardiovascular system associated with abiraterone.

**Fig. 6.**
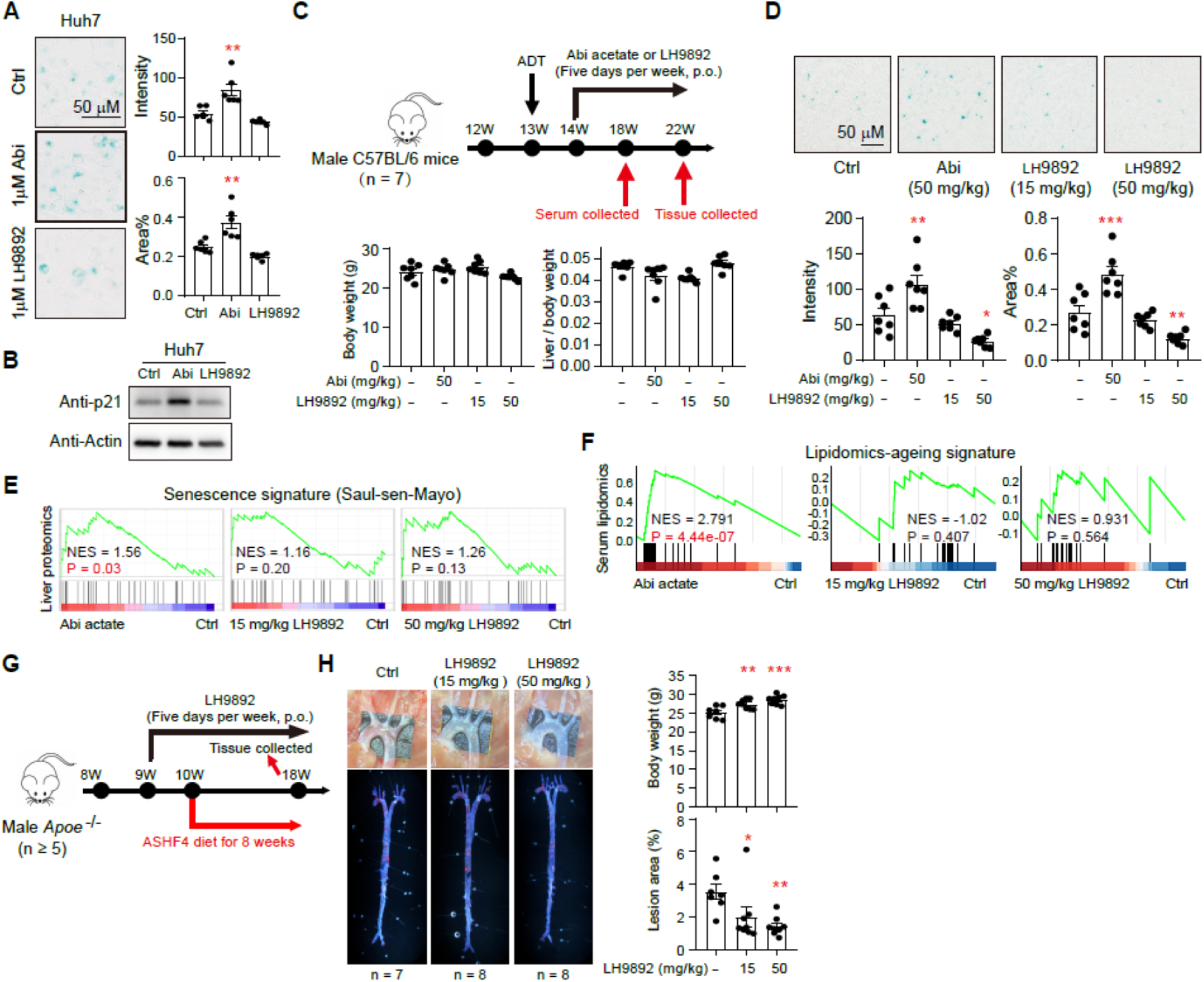
Effect of LH9892 on systemic metabolism. **A**, SA-β-gal staining in Huh7 cells treated with abiraterone (Abi, 1 µM) or LH9892 (1 µM) for 24 hrs. Representative images are shown (n = 6 images per group). **B**, p21 expression in Huh7 cells treated with abiraterone (1 µM) or LH9892 (1 µM) for 48 hrs. **C**, Mice body weight and liver weight after abiraterone (50 mg/kg/day, per os) or LH9892 (15 or 50 mg/kg/day, per os) treatment (n = 7 per group). **D**, Quantification of SA-β-gal-positive area per field in liver sections (n = 7 per group) from male mice treated with abiraterone acetate or LH9892. **E**, GSEA of the Saul-sen-Mayo senescence signature on proteomics of liver tissues from mice treated with abiraterone or LH9892 (n = 4 per group except ctrl group with 3 mice). **F**, Enrichment analysis on lipidomics-ageing signature with serum samples from abiraterone or LH9892-treated mouse (n = 7 per group). **G**, A schematic showing evaluation of atherosclerosis procedure in male *Apoe*^⁻/⁻^ mice. Mice were treated with or without LH9892 (15 or 50 mg/kg/day) via oral gavage 5 days per week. Mice were fed with ASHF4 diet for 8 weeks. **H**, Representative images of atherosclerotic plaques in the aorta and Oil Red O staining of en face aorta from male *Apoe*^-/-^ mice treated with or without LH9892 (Ctrl group, n = 7; LH9892 treated mice, n = 8 per group). Data are shown as the mean ± SEM. *, ** and *** denoted P < 0.05, P < 0.01 and P < 0.001, respectively.

## Discussion

Cancer therapy has long prioritized cancer cell eradication or control over the long-term systemic toxicity of therapeutic agents (1, 2). The ‘tolerable’ drug-induced side effects reflect compromised organismal homeostasis, which is intolerable latent threats to wellbeing, particularly for prostate cancer which has much longer treatment durations. Here we integrated analyses of clinical samples and metabolomics to decipher abiraterone-induced systemic health detriments and the underlying mechanisms. Leveraging structural biology and structure-activity relationship studies, we developed LifeHug-9892 to mitigate the impairment to organismal health while preserving antitumor activity. Our findings lay a foundation for the development of third-generation therapy drugs for prostate cancer.

Hepatic and cardiovascular disorders are well-documented adverse effects of abiraterone (6, 8, 9, 20). As abiraterone is administered earlier in the disease course, its differential benefits in prolonging progression-free survival and overall survival become increasingly pronounced. Concomitantly, the risks of hypertension and hepatic disorders rise with prolonged abiraterone administration (6, 8, 9, 20). These observations underscore that these are not merely ‘tolerable’ side effects but potential contributors to life-threatening risks in prostate cancer patients. Mitigating or preventing abiraterone-induced interference with organismal metabolic homeostasis is critical to accommodate the extended treatment duration of abiraterone.

Unlike the side effects of ADT, which primarily arise from direct or indirect perturbations of androgen physiology and thus remain challenging to mitigate — abiraterone-induced metabolic dysregulation is androgen-independent (21). In mice, which lack adrenal CYP17A1, abiraterone minimally disrupts androgen homeostasis yet recapitulates the metabolic changes observed in humans. This creates opportunities for drug optimization to reduce its systemic toxic effects while preserving its antitumor activity. Our time-series metabolomic analysis of patient cohorts reveals that abiraterone induces extensive and persistent metabolomic perturbations, particularly a marked elevation in desmosterol levels. Desmosterol exhibits higher cellular efflux efficiency than cholesterol, a property enhancing its capacity for inter-organ crosstalk (22). Notably, elevated desmosterol has been implicated in the pathogenesis of cardiovascular disease, non-alcoholic steatohepatitis, hepatitis C virus infection, and other pathological conditions (23–27). While decreased desmosterol levels in blood or brain serve as an ageing marker (28, 29), these physiological alterations are mild and serve as an indicator for endogenous cholesterol synthesis. In contrast, abiraterone-induced desmosterol accumulation reaches over 10-fold upregulation in both mice and patients. Dosage is critical to metabolite function (30). Consistently, marked desmosterol elevation via DHCR24 knockdown has been shown to induce HUVEC senescence (31). We also found enhanced SA-β-gal activity in liver cells and tissues following abiraterone treatment. The liver is prone to senescence, a process that is particularly difficult to reverse (32, 33), which may mechanistically explain abiraterone-induced hepatic dysfunction. Considering the different effect of 5α-Abis and 5β-Abis on DHCR24 (Fig. 2E), patients with higher SRD5A activity might be more vulnerable to the desmosterol-related adverse events (34).

Our previous studies revealed that abiraterone is recognized by steroidogenic enzymes, including 3βHSD1 and SRD5A, to generate D4A and other bioactive metabolites with diverse pathological effects (17, 18). Given its steroidal scaffold, it is not surprising for abiraterone to interact with other metabolic enzymes. Direct binding and inhibition of DHCR24 by abiraterone is consistent with the rapid and robust desmosterol elevation observed in mice within 12 hours (hrs) of abiraterone administration. Such pronounced and persistent desmosterol accumulation in abiraterone-treated patients is exceptionally rare, indicating that systemic metabolic homeostasis is unable to adapt to resolve abiraterone-induced cholesterol metabolism dysregulation. Notably, while abiraterone inhibits DHCR24, it ultimately upregulates of both desmosterol (DHCR24 substrate) and cholesterol (DHCR24 product). Mechanistically, abiraterone binding to DHCR24 not only inhibits DHCR24 enzymatic activity but also protects DHCR24 from AMFR-mediated degradation. Accumulated desmosterol fuels cholesterol synthesis when serum abiraterone concentrations decline—driven by either metabolic rhythmicity or long-term abiraterone treatment (1, 35). This creates a unique metabolic scenario where both the substrate (desmosterol) and product (cholesterol) accumulate after DHCR24 inhibition, exacerbating abiraterone-induced metabolic dysregulation and increasing cardiovascular diseases risk.

Statins or ezetimibe may mitigate abiraterone-induced metabolic dysregulation, but their potential side effects must be considered (36). Ideally, the development of third-generation ARPIs with attenuated systemic toxicity is warranted. Second-generation of ARPIs have focused primarily on tumor cells, enhancing prostate cancer inhibition by improving CYP17A1 or AR inhibitory efficacy. The design of third-generation ARPIs should adopt a patient-centric paradigm, prioritizing not only tumor control but also preservation of systemic health and metabolic homeostasis. With advances of artificial intelligence (AI), identifying off-target drug effects is expected to move beyond labor-intensive biochemical screening. Additionally, the evolution of metabolomics and other multi-omics technologies enables comprehensive assessment of global physiological perturbations at the patient level, providing insights into systemic metabolic dynamics beyond tumor-specific changes. These technological and conceptual advances lay a solid foundation for the development of third-generation ARPIs with improved safety profiles and sustained anticancer efficacy.

In summary, our findings demonstrate that abiraterone inhibits DHCR24, leading to systemic metabolic dysregulation and imposing potential risks to organismal health. LH9892, targeting the intolerable tolerance, is designed to effectively inhibit prostate cancer progression while maintaining systemic metabolic homeostasis.

## Materials and Methods

### Ethical statements

All studies involving human participants were conducted in accordance with the principles of the Declaration of Helsinki and approved by the Ethics Committee of Tongji Hospital, Shanghai, China (2018-LCYJ-003 and 2019-007). Written informed consent was obtained from all participants.

Animal experiments were performed in compliance with the guidelines of the Institutional Animal Care and Use Committee (IACUC) of the Institute of Biochemistry and Cell Biology, Shanghai Institutes for Biological Sciences, Chinese Academy of Sciences. All procedures were designed to minimize suffering, and tumor sizes did not exceed the limits approved by the IACUC.

### Materials and cell lines

LNCaP, PC-3 and HEK293T cells were obtained from the American Type Culture Collection (ATCC, Manassas, VA, USA). VCaP cells were kindly provided by Dr. Jun Qin (Shanghai Institute of Nutrition and Health, Chinese Academy of Sciences, Shanghai, China). Huh7, NIH/3T3, and THP-1 cells were acquired from the Cell Bank/ Stem Cell Bank of the Chinese Academy of Sciences. Human umbilical vein endothelial cells (HUVECs) were obtained from the Shanghai Huzhen Industrial Co., Ltd (HZ-H205). Cell lines were authenticated by Hybribio (Guangzhou, China) and determined to be mycoplasma free with primers 5’-GGGAGCAAACAGGATTAGATACCCT -3’ and 5’-TGCACCATCTGTCACTCTGTTAACC TC -3’.

LNCaP, THP-1, and PC3 cells were maintained in RPMI-1640 medium supplemented with 10% fetal bovine serum (FBS). Huh7, NIH/3T3, HEK293T, VCaP, and VCaP-CYP17A1 cells were grown in DMEM with 10% FBS. Puromycin (1 µg/ml) was used for VCaP-CYP17A1 selection. HUVECs were cultured in endothelial cell medium (ECM) containing 5% FBS and 1% endothelial cell growth supplement (ECGS). All cell lines were cultured at 37 °C in a humidified atmosphere with 5% CO₂. To induce differentiation, THP-1 cells were treated with 100 ng/mL phorbol 12-myristate 13-acetate (PMA) in differentiation medium (DMEM containing 10% FBS) for 48 hrs after seeding.

Primary antibodies used for immunoblots were as follows: rabbit monoclonal antibody against p21 (Abcam, Cambridge, UK; ab109520, 1:1000), rabbit polyclonal antibody against p21 (Proteintech, Rosemont, IL, USA; 28248-1-AP, 1:1000), rabbit monoclonal antibody against β-actin (ABclonal, China; AC026, 1:10000), rabbit polyclonal antibody against CD36 (Proteintech, Rosemont, IL, USA; 18836-1-AP, 1:1000), rabbit polyclonal antibody against DHCR24 (Proteintech, Rosemont, IL, USA; 10471-1-AP, 1:1000), rabbit polyclonal antibody against AMFR (Cell Signaling Technology, Danvers, MA, USA; 9590S, 1:1000), mouse monoclonal antibody against HA (Abmart, China; M20003L, 1:1000), rabbit polyclonal antibody against Flag (Proteintech, Rosemont, IL, USA; 20543-1-AP), mouse monoclonal antibody against DHCR24 (Santa Cruz Biotechnology, Dallas, Texas, USA; sc398938, 1:000), rabbit polyclonal antibody against MYC (Sigma Aldrich, St. Louis, MO, USA; 06-549, 1:1000).

Primary antibodies used for immunoprecipitation were as follows: rabbit monoclonal antibody against HA (Cell Signaling Technology, Danvers, MA, USA; 3724T, 1:100), rabbit monoclonal antibody against DHCR24 (Cell Signaling Technology, Danvers, MA, USA; 2033S, 1:100), mouse monoclonal antibody against Flag (Sigma Aldrich, St. Louis, MO, USA; F1804, 1:100).

Antibodies used for immunofluorescence were as follows: mouse monoclonal antibody against CD68 (Novus Biologicals, Centennial, CO, USA; NB100-683, 1:200), rabbit monoclonal antibody against α-Smooth Muscle Actin (SMA) (Cell Signaling Technology, Danvers, MA, USA; 19245s, 1:400), Alexa Fluor^TM^488 donkey anti-mouse IgG (H+L) (Invitrogen, Waltham, MA, USA; A21202, 1:1000), Alexa Fluor^TM^594 donkey anti-rabbit IgG (H+L) (Invitrogen, Waltham, MA, USA; A21207, 1:1000).

### Patient cohort

Clinical specimens from 15 patients with metastatic hormone-sensitive prostate cancer (mHSPC) were retrospectively collected from the biobank of Shanghai Tongji hospital. These patients were treated with first-line abiraterone acetate (1000 mg/day) plus prednisone (5 mg, twice daily) combined with androgen deprivation therapy (ADT), and have sequential serum samples at baseline as well as at 1, 2, 3, and more than 7 months post-treatment. None of these patients present with hepatic dysfunction, unstable cardiac disease, or hyperlipidemia at the baseline of treatment. Written informed consent was obtained from all participants, and the study protocol was approved by the Institutional Review Board of Shanghai Tongji Hospital (2018-LCYJ-003 and 2019-007).

### Serum collection

Blood samples were obtained from both prostate cancer patients and experimental mice. Patient blood was collected via venipuncture into standard coagulation vacuum tubes containing nano-scale microsphere silica as a coagulant, and centrifuged at 3,000 rpm for 3 minutes at 4°C. Mouse blood was collected via retro-orbital puncture, and centrifuged at 9,000 rpm for 15 minutes at 4°C. The resulting supernatant (serum) was aliquoted and stored at −80°C until further use.

### Targeted lipidomics

Lipid extraction and detection were performed based on previously described protocols with minor modifications (37, 38). Serum samples were mixed with MTBE for lipid extraction. Ultra-high performance liquid chromatography-mass spectrometry (UPLC-MS) was conducted using a Shimadzu chromatography system alongside a SCIEX 6500 mass spectrometry system for targeted lipidomics analysis with different methods. Samples were analyzed with an Agilent Zorbax Eclipse Plus C18 (2.1 x 100 mm, 1.8 μm), Waters BEH HILIC (2.1 x 100 mm, 1.7 μm), using 1 μL sample injections, and a Phenomenex Kinetex C18 (2.1 x 100 mm, 2.6 μm) using 2 μL sample injections. Gradient elution for the EC18 column was performed using a mobile phase consisting of water/methanol/acetonitrile (1:1:1, v/v/v, containing 0.1% formic acid and 1 mM ammonium acetate) as phase A and isopropanol/acetonitrile (90:10, v/v, containing 0.05% formic acid and 10 mM ammonium acetate) as phase B. For the HILIC column, the mobile phases were water/acetonitrile (5:95, v/v, containing 10 mM ammonium acetate) as phase A and water/acetonitrile (1:1, v/v, containing 10 mM ammonium acetate) as phase B. For the C18 column, gradient elution utilized water/methanol/acetonitrile (1:1:1, v/v/v, containing 7 mM ammonium acetate) as phase A and isopropanol (containing 7 mM ammonium acetate) as phase B. Data were acquired through MRM scanning. All raw UHPLC-MS data were acquired by Analyst software (SCIEX, version 1.6). Raw data files were then imported to the SCIEX OS (SCIEX, version 3.0) for the peak extraction, alignment, and peak area calculation.

### Sterols

Quantitative method of sterol metabolites was conducted as reported previously (39). In brief, 50 μL serum sample was added to 50 μL of an internal standard mixture. Then, 500 μL of pre-cooled methanol (−20°C) was added, vortexed, and centrifuged at 12,000 rpm at 4°C for 10 min to obtain the supernatant. This extraction procedure was repeated once more and the two supernatants from each sample were pooled and dried using nitrogen gas. The residue was re-dissolved in 50 μL of acetonitrile. Then, the nitrogen-dried residue of the working solution (10 μL) and biological sample extracts were added to 100 μL pyridine solution of PA, MNBA, and DMPA, respectively. The mixtures were heated at 80°C for 60 min, followed by the addition of pure water (200 μL) and extraction with MTBE (1.5 mL). These MTBE extracts were dried with nitrogen gas, and the resultant residues were reconstituted in 100 μL acetonitrile for the UHPLC-MS/MS analysis of steroid hormones and hydroxysterols. Subsequently, a 10 μL aliquot of this reconstitution solution was diluted with 200 μL acetonitrile for the targeted analysis of steroid metabolites in the Kandutsch–Russell pathway and Bloch pathway. Quantification was performed on an LC-30AD UHPLC system (Shimadzu Technologies, Japan) hyphenated with a 6500 plus Qtrap mass spectrometer (SCIEX, Framingham, MA, USA) with an electrospray ionization (ESI) source. A Zorbax Eclipse Plus C18 column (100 × 2.1 mm, 1.8 μm; Agilent Technologies, Santa Clara, CA, USA) was used with water and a mixture of acetonitrile and isopropanol (10:1, v/v) containing 0.1% formic acid as mobile phases A and B, respectively.

### Untargeted metabolomics

20 µL serum samples were processed using a modified methanol/MTBE liquid-liquid extraction. Protein precipitation and metabolite extraction were initiated by adding 100 µL of ice-cold methanol. The mixture was vigorously vortexed for 30 seconds and then centrifuged at 14,000 × g and 4°C for 10 min. Subsequently, 90 µL of the supernatant was carefully transferred to a new tube for a biphasic liquid-liquid extraction. This was achieved by adding 250 µL of pre-cooled MTBE, vortexing for 30 seconds, followed by the addition of 47.5 µL of pure water and another 30 seconds of vortexing to induce phase separation. The mixture was then centrifuged at 14,000 × g and 4°C for 5 min. Following centrifugation, approximately 280 µL of the upper organic layer (MTBE phase) was carefully aspirated and discarded. The lower aqueous layer, containing the extracted polar metabolites, was collected. An 80 µL aliquot of this aqueous phase was transferred into a glass insert placed within a chromatographic vial for immediate LC-MS/MS analysis. Chromatographic separation was performed on a Waters ACQUITY UPLC HSS T3 (100 × 2.1 mm, 1.8 µm) and Waters ACQUITY UPLC BEH Amide (100 × 2.1 mm, 1.7 µm) using Waters ACQUITY UPLC system. The mobile phases of T3 column consisted of water with 0.1% formic acid (A) and acetonitrile with 0.1% formic acid (B) at a flow rate of 0.3 mL/min with a gradient elution. The mobile phases of Amide column consisted of 5:95 (v/v) acetonitrile/water containing 0.3% (v/v) formic acid and 10 mM ammonium formate (A) and 5:95 (v/v) water/acetonitrile containing 0.3% (v/v) formic acid and 10 mM ammonium formate (B) at a flow rate of 0.3 mL/min with a gradient elution. Mass spectrometry analysis was conducted on a Waters Xevo G2-XS QTOF with ESI source operating in both positive and negative modes. Key MS parameters were: capillary voltage, 2.5 kV (positive) and 2.0 kV (negative); source temperature, 120°C; desolvation temperature, 500°C; cone and desolvation gas flows at 150 L/hr and 800 L/hr, respectively.

### Real-time quantitative PCR (RT-qPCR)

For direct cDNA synthesis from cultured cells, the Cell to cDNA Kit (EZBioscience, China; B0015) was employed. Total RNA from mouse tissues was isolated using TRIzol® Reagent (Invitrogen, Waltham, MA, USA; 15596018CN). cDNA was synthesized using HiScript II Q RT SuperMix for qPCR (+gDNA wiper) (Vazyme, China; R223-01) according to the manufacturer’ s instructions. Quantitative PCR was performed with 2 x SYBR Green qPCR Master Mix (EZBioscience, China; A0001) on a Roche LightCycler® 96 system. Each reaction was run in technical triplicates.

### In vivo ubiquitination assay

In vivo ubiquitination was performed as described previously (2). Briefly, following transfection or drugs treatment, cells were collected and lysed in RIPA buffer supplemented with 1.0% protease inhibitor cocktail. Cell lysates were then sonicated and centrifuged. Following 6 hrs incubation of indicated beads with cell supernatant, the beads were washed 4 times with RIPA buffer for western blot assay.

### Surface plasmon resonance

Surface plasmon resonance (SPR) analysis was performed on a Biacore 8K system (Cytiva, Marlborough, MA, USA) in a running buffer containing 25 mM HEPES (pH 7.4), 150 mM NaCl, 5% (v/v) DMSO, 0.001% (w/v) LMNG, and 0.0002% (w/v) CHS. The assay was conducted at 25°C with a flow rate of 30 µL/min. Purified DHCR24 protein was diluted in sodium acetate buffer (pH 4.5) and immobilized on the Series S CM5 sensor chips (Cytiva, Marlborough, MA, USA) using 1-ethyl-3-(3-dimethylaminopropyl) carbodiimide hydrochloride/N-hydroxysuccinimide (EDC/NHS) coupling reagent, yielding a resonance response of approximately 20,000 response units (RUs). A concentration gradient of different compounds was injected over the sensor chip surface. The results were analyzed with a steady state affinity binding model, and the data were fitted and evaluated using the Biacore Insight Evaluation Software (Version 3.0.12).

### Immunofluorescence

Tissue samples were dissected in PBS and fixed overnight at 4°C in 4% paraformaldehyde (PFA). Samples were then embedded in optimal cutting temperature (OCT) compound and cryo-sectioned at a thickness of 10 μm. For immunofluorescence, sections were rehydrated in PBS and post-fixed with 4% PFA. Permeabilization was performed using 0.2% Triton X-100 in PBS, followed by blocking with 5% normal goat serum in PBS. Sections were incubated with primary antibodies overnight and species-appropriate fluorescent secondary antibodies. Nuclei were stained with 4′,6-diamidino-2-phenylindole (DAPI). Images were acquired using an Olympus BX53 fluorescence microscope.

### SA-β-gal staining

Cellular senescence was assessed by senescence-associated β-galactosidase (SA-β-gal) activity using the Senescence β-Galactosidase Staining Kit (Beyotime, China; C0602) following the manufacturer’s protocol. In brief, cells or frozen tissue sections were fixed with fixation solution at room temperature. The staining working solution was prepared by combining SA-β-galactosidase staining solution A, B, C, and X-Gal solution according to the kit instructions. Samples were covered with the working solution and incubated in a humidified dark environment at 37°C without CO₂. Images were captured using an Olympus microscope (BX53 or IX73) under bright-field settings. Quantification of SA-β-gal-positive cells was performed using ImageJ software (v1.53).

### Oil Red O staining

Adherent cells or cryo-sectioned liver tissues (10 μm) were fixed in 4% paraformaldehyde (PFA) for 15 minutes at room temperature. Fixed samples were incubated in freshly prepared Oil Red O working solution for 30 minutes at room temperature. Following incubation, samples were differentiated in 60% isopropanol to remove non-specific staining, and washed thoroughly with PBS. Nuclei were counterstained with hematoxylin for 1 minute. Finally, samples were imaged using an Olympus microscope (BX53 or IX73). Lipid droplet area was quantified using ImageJ software (v1.53).

### Cell proliferation assay

Cell proliferation was assessed using the Cell Counting Kit-8 (Beyotime, China; C0040) according to the manufacturer’s instructions. Cells were seeded in 96-well plates and allowed to adhere for 24 hrs. Compounds were then added at the indicated concentrations. At each time point, 10 μl of CCK-8 reagent was added to each well, and incubated with cells. Absorbance was measured at 450 nm with a reference wavelength of 600 nm using a BioTek microplate reader (BioTek Instruments, Winooski, VT, USA). All experiments were performed with at least three technical replicates and repeated in three independent biological replicates.

### Cell death assay

Cell death was analyzed using the Annexin V-PE/DAPI Apoptosis Detection Kit (APExBIO, Houston, TX, USA; K2282). Cells were seeded in 6-well plates and treated with the indicated compounds for 72 hrs. After treatment, cells were collected in 1× binding buffer for staining using Annexin V-PE and DAPI. Samples were analyzed immediately using a BD LSRFortessa flow cytometer (BD Biosciences, San Jose, CA, USA). A minimum of 10,000 events were recorded per sample, and data were processed using FlowJo software (v10.8.1).

### Bulk RNA sequencing

Total RNA was isolated from each sample using TRIzol reagent. Library preparation was carried out with the VAHTSTM mRNA-seq V3 Library Prep Kit for Illumina (NR611) according to the manufacturer’s protocol. Briefly, 1 μg of total RNA was used for mRNA enrichment and fragmentation. The purified mRNA was then subjected to first-and second-strand cDNA synthesis. Double-stranded cDNA was ligated to Illumina-compatible sequencing adapters (VAHTSTM RNA Adapters set3–set6, N809/N810/N811/N812) and amplified via PCR. Final libraries were quantified using a Qubit Fluorometer (Invitrogen, Waltham, MA, USA) and assessed for quality with an Agilent 2100 BioAnalyzer. Sequencing was performed on an Illumina HiSeq platform with a paired-end 150 bp read configuration. Raw sequencing data were quality-controlled with fastp (v0.23.4). Paired-end reads were aligned to the human reference genome GRCh38.91 using STAR (v2.7.10b). Gene-level quantification was performed with featureCounts (v2.0.6). Read counts were normalized and differential gene expression analysis was conducted using the DESeq2 package (v1.36.0) in R. Biological replicates were included for each experimental condition.

### Preparation of conditioned medium from HUVECs

HUVECs were cultured in endothelial cell medium (ECM) and treated with desmosterol (10, 20, or 50 μM) or cholesterol (10, 20, or 50 μM) for 72 hrs. Conditioned medium was collected, centrifuged, and filtered through a 0.22 μm membrane. Aliquots were stored at −80°C until further use.

### MTBE-based extraction and functional analysis of conditioned medium fractions

To characterize bioactive soluble mediators secreted by HUVECs, conditioned medium (CM) was subjected to biphasic extraction using methyl tert-butyl ether (MTBE). CM was combined with pre-chilled MTBE at a ratio of 1:2.5 (v/v), vigorously vortexed, and centrifuged at 12,000 g for 3 minutes to achieve phase separation. The upper organic and lower aqueous phases were carefully isolated. The organic phase was evaporated under nitrogen and reconstituted in anhydrous ethanol. The aqueous phase was lyophilized and redissolved in phosphate-buffered saline (PBS). Both fractions were normalized to volumes equivalent to their original proportions in CM.

For functional assessment, THP-1 cells were treated with either the ethanol-reconstituted organic fraction or the PBS-reconstituted aqueous fraction. Vehicle controls received equivalent volumes of ethanol or PBS. After 12 hrs of treatment, transcriptional changes in *CD36*, *ABCA1* and *RPLP0* were quantified using RT-qPCR. All experiments included at least three biological replicates.

### Thermal inactivation and functional profiling of conditioned medium

To evaluate the contribution of thermolabile protein factors to bioactivity, HUVEC-derived CM was heat-inactivated at 100°C for 10 minutes using a dry bath heater.

Samples were then centrifuged at 12,000 g for 10 minutes to remove insoluble aggregates. The clarified supernatant was applied to THP-1 monocytes. After 12 hrs of stimulation, *CD36* expression was analyzed via RT-qPCR using *RPLP0* as a reference gene.

### Protease-based degradation and functional validation of conditioned medium

To further assess protein-dependent bioactivity, CM was treated with protease K (1 mg/mL) and incubated at 37°C for 2 hrs to digest protein components. Protease activity was terminated by heating at 100°C for 10 minutes, followed by centrifugation at 12,000 g for 10 minutes to remove precipitates. The resulting supernatant was applied to THP-1 cells. Parallel treatments with native CM, vehicle, and protease K-treated non-heated CM were included as controls. After 12 hrs, *CD36* mRNA levels were measured by RT-qPCR and normalized to *RPLP0* expression. All experiments were performed in triplicate.

### Androgen metabolism tracing assay

Tracing assay with HPLC analysis was performed as previously described with modifications. Briefly, cells were seeded in 24-well plates. After 24 hrs, cells were treated with the indicated compounds and 1 × 10^5^ cpm/well of [³H]-pregnenolone, [³H]-dehydroepiandrosterone and [³H]-androstenedione (PerkinElmer, Waltham, MA, USA) at 37°C. At indicated time point, 200 μl of culture medium was collected for metabolite analysis.

Samples were incubated with β-glucuronidase (Novoprotein Scientific Inc., China) at 37°C for 2 hrs to hydrolyze conjugated metabolites. Metabolites were extracted with a 1:1 mixture of ethyl acetate and isooctane, and the organic phase was dried using a freeze dryer. Dried extracts were dissolved in 100 μl of 50% methanol and injected into an Acquity Arc HPLC System (Waters, Milford, MA, USA). Separation was performed on a CORTECS C18 reversed-phase column (Waters, Ireland) maintained at 40°C, using a methanol/water gradient. Radio-labeled metabolites were detected in real time using a β-RAM model 3 in-line radioactivity detector (LabLogic Systems). A mixture of [³H]-labeled standards (DHEA, progesterone, pregnenolone) was used for metabolite identification. Quantification was based on the area under the curve (AUC) for each metabolite peak.

### Mouse models

Male mice were used throughout this study. Wild-type C57BL/6J mice (8 or 12 weeks old) were purchased from Lingchang Biotech (Shanghai, China). B-NDG mice (B: Biocytogen; N: NOD background; D: DNAPK (Prkdc) null; G: IL2rgknockout; 4-6 weeks old) were obtained from Biocytogen (Beijing, China). *Apoe*^‒/‒^ mice on a C57BL/6 background for atherosclerosis were acquired from GemPharmatech Co., Ltd (Nanjing, China). Mice were housed under specific pathogen-free (SPF) conditions in the animal facility of the Institute of Biochemistry and Cell Biology. For each experiment, age-matched mice were randomly assigned to control or experimental groups.

### Mice metabolism model

Eight-week-old male C57BL/6J wild-type mice were randomly assigned to either a control group or an abiraterone acetate (Abi) treatment group. The Abi group received daily intraperitoneal injections of abiraterone acetate at a dose of 0.15 mmol/kg/day for seven consecutive days, while the control group received vehicle injections on the same schedule. Serum samples were collected at baseline, day 1, day 4, day 7, and on the first day after treatment cessation (day 8) for subsequent lipidomic and steroidomic profiling.

To assess the metabolic effects of abiraterone’s metabolites, eight-week-old male C57BL/6J wild-type mice were randomly assigned to five groups: vehicle, abiraterone acetate (0.15 mmol/kg/day), D4A (0.15 mmol/kg/day), 5α-abiraterone (0.15 mmol/kg/day), or 5β-abiraterone (0.15 mmol/kg/day). Compounds were administered once daily via intraperitoneal injection over four consecutive days. Serum samples were collected at baseline, day 1, and day 4 for subsequent lipidomic and steroidomic profiling.

In a separate experiment designed to evaluate LH9892’s effect on metabolism, male mice were divided into three groups: vehicle, abiraterone acetate (0.15 mmol/kg/day), and LH9892 (0.15 mmol/kg/day). Treatments were delivered daily via intraperitoneal injection for one week. Serum was similarly collected at multiple time points for lipidomic and steroidomic analyses. At the experimental endpoint, mice were euthanized under anesthesia and transcardially perfused with PBS. The heart, liver, brain, adrenal glands, prostate, adipose tissue, kidney, lung, and spleen were harvested and snap-frozen at –80°C for subsequent drug metabolism studies.

### Xenograft studies

VCaP-CYP17A1 cells (1 x 10⁷) were resuspended in Matrigel (Corning, Corning, NY, USA; 354234) and implanted subcutaneously into the right flank of intact male mice. When tumor volumes reached 200-300 mm³, mice were castrated and implanted subcutaneously with sustained-release pregnenolone pellets. Mice were then randomized into treatment groups when tumors reached approximately 200 mm³. Animals were treated daily via oral gavage (p.o.) for 3 weeks with the following: vehicle, abiraterone acetate (50 mg/kg/day) or LH9892 (15 or 50 mg/kg/day). Tumor dimensions were measured every two days using calipers, and volumes were calculated as V = L × W² × 0.52 (L: length; W: width). The humane endpoint for survival analysis was defined as a tumor volume exceeding 1,500 mm³ or the manifestation of ethical euthanasia criteria as assessed by a veterinarian. All cells used for in vivo implantation were confirmed mycoplasma-free.

### Abiraterone induced senescence mouse model

Three-month-old male C57BL/6J mice were randomly assigned to control or abiraterone acetate (Abi) treatment groups. The Abi group received subcutaneous implantation of sustained-release abiraterone acetate pellets, which the control group underwent a sham implantation procedure. At the endpoint, mice were euthanized under anesthesia and transcardially perfused with PBS. Liver, heart, spleen, kidney, lung, brain, and muscle tissues were dissected and fixed in 4% PFA for subsequent histological evaluation.

### Therapeutic intervention mouse model

Three-month-old male C57BL/6J wild-type mice were randomly assigned to four treatment groups: mock, abiraterone acetate (50 mg/kg/day), LH9892 (15 mg/kg/day), or LH9892 (50 mg/kg/day). Treatments were administered via oral gavage on a 5-days-on/2-days-off schedule for 2 months. Serum samples were collected after one month of treatment. At the endpoint, mice were euthanized and processed as described above.

### ASHF4 diet-induced mouse atherosclerosis model

Male *Apoe*^-/-^ mice (8 weeks old) were randomly assigned to control group or abiraterone acetate-treated groups. The abiraterone group received subcutaneous implantation of sustained-release abiraterone acetate pellets, which the control group underwent a sham implantation procedure. After one week, all mice were fed an ASHF4 diet (containing 40 kcal% fat, 1.25% cholesterol, 0.5% sodium cholate) for 5 weeks. At 15 weeks of age, mice were euthanized under anesthesia and transcardially perfused with PBS. The heart and entire arterial tree were carefully dissected and fixed in 4% paraformaldehyde (PFA) for subsequent histological analysis.

### Normal diet-induced long-term atherosclerosis model

Male *Apoe*^-/-^ mice (8 weeks old) were randomly divided into control group or abiraterone-treated groups. The abiraterone group received subcutaneous implantation of sustained-release abiraterone acetate pellets, which the control group underwent a sham implantation procedure. All mice were maintained on a standard normal diet for 6 months. At the endpoint, mice were euthanized and processed as described above.

### Atherosclerotic lesion analysis

For Oil Red O staining of the entire aorta, the aorta samples were fixed with 4% polyformaldehyde overnight and washed with phosphate buffered saline (PBS). The aorta samples were then cut open longitudinally and stained with 0.3% Oil Red O (Sigma Aldrich, St. Louis, MO, USA ; 1320-06-5) for 20 min, followed by 60% isopropanol for 5 min. The Oil Red O-positive areas of each aorta sample were captured by stereo microscope and evaluated using the ImageJ software.

For Oil Red O staining of the aortic root, the frozen sections subjected to Oil Red O as previous described, rinsed with water, and then stained with hematoxylin for 2 min, finally sealed with glycerol. The Oil Red O-positive areas of sections were captured by Olympus microscope (BX53F) and evaluated using the ImageJ software.

For Immunofluorescent staining of the aortic intact tricuspid valve, the frozen sections were permeabilized and blocked with 0.3% Triton X-100 and 5% bovine serum albumin (BSA) at room temperature for 1h, and incubated with mouse anti-CD68 and rabbit anti-*α*-smooth muscle actin (SMA) overnight at 4°C. After washing with PBS, the sections were incubated with the Alexa FluorTM488 donkey anti-mouse IgG(H+L) and Alexa FluorTM594 donkey anti-rabbit IgG(H+L) for 1 h in dark. The sections were then stained with DAPI for 10 min at room temperature. Fluorescent images were captured using a fluorescence microscope (Keyence, IL, USA; BZ-X810), and the fluorescent area was quantified using ImageJ software.

### Therapeutic intervention atherosclerosis model

Male *Apoe*^-/-^ mice (8 weeks old) were randomly assigned to three treatment groups: vehicle, LH9892 (15 mg/kg/day), or LH9892 (50 mg/kg/day). Animals were treated via oral gavage on a 5-days-on/2-days-off schedule for 9 weeks. Throughout the experimental period, mice were maintained on an ASHF4 diet (40 kcal% fat, 1.25% cholesterol, 0.5% sodium cholate). At the endpoint, mice were euthanized and processed as described above.

### Boltz-2 sequence-based virtual screening and affinity prediction

Potential abiraterone targets were evaluated using Boltz-2, which co-predicts protein-ligand complex structures and binding affinity from amino-acid sequences and small-molecule SMILES. For each candidate protein, the full sequence was supplied; when a catalytic cofactor or metal ion was known, the corresponding small molecule/ion was included as an additional SMILES in the receptor context. The abiraterone ligand (SMILES retrieved from ChEMBL) was used for all runs. Boltz-2 was executed with default parameters uniformly across targets, and for each protein the top-scoring pose and the model-reported Affinity (nM) were retained. For visualization and ranking, affinities were transformed to *p*Affinity = 3 − log_10_(Affinity [nM]), and proteins were ordered from highest to lowest pAffinity.

### Statistics

Bar plots show quantitative data as mean ± standard error of the mean (SEM); violin plots show median, 25^th^ percentile, and 75^th^ percentile with the outer shape showing the kernel density estimation of the data distribution. For statistical analysis, statistical methods were chosen according to experimental design and data properties.

Comparisons between two groups: For two independent groups, if data were normal and variances equal, then unpaired two-tailed Student’s *t*-test was applied; if normal but variances unequal, then unpaired two-tailed *t*-test with Welch’s correction was applied; otherwise (non-normal), the Mann-Whitney test was applied. For two paired groups, if normal, then paired two-tailed *t*-test was applied; otherwise, Wilcoxon matched-pairs signed rank test was applied.

Comparisons among three or more groups: For multiple independent groups, if normal with equal variances, then one-way ANOVA followed by Dunnett’s post hoc test was applied; if normal with unequal variances, then Welch’s ANOVA followed by Dunnett’s T3 post hoc test was applied; otherwise, the Kruskal-Wallis test followed by Dunn’s post hoc test was applied. For repeated measures, if normal and spherical, then repeated-measures (RM) ANOVA followed by Holm-Sidak post hoc test was applied; if normal but not spherical, then RM one-way ANOVA with Greenhouse-Geisser correction and Holm-Sidak post hoc test was applied; otherwise, Friedman test followed by Dunn’s post hoc test was applied.

Comparisons involving two independent factors: Two-way ANOVA was performed. For post hoc comparisons among multiple groups, Dunnett’s test (for comparisons against control) or the Holm-Šidák test (for pairwise comparisons) was applied, as specified in the Figure legends.

Specific to the lipidomics dataset, paired comparisons were assessed using the Wilcoxon matched-pairs signed-rank test, and the Storey method was applied to control the false discovery rate (FDR) for multiple hypothesis testing. Trend analysis was carried out with Short Time-series Expression Miner (STEM), and visualizations were created in R. Graph generation and Figure assembly were done using GraphPad Prism 8.0 and Adobe Illustrator, respectively.

For all figures, *** indicates *P* < 0.001, ** indicates *P* < 0.01, and * indicates *P* < 0.05.

## Acknowledgements

We thank the staff members (Pengyu Wang) of Bio-Med Big Data Center/ Shanghai Institute of Nutrition and Health (SINH) and staff members (Qingjie Xiao) of the Shanghai Synchrotron Radiation Facility (SSRF) beamline BL18U1 for providing invaluable technical support and assistance in data collection and analysis. We acknowledge financial supports from the National Key R&D program of China (2023YFC3402700), the Strategic Priority Research Program of the Chinese Academy of Sciences (XDB0990000), National Key R&D program of China (2023YFC3404700; 2022YFC3400700, 2022YFA0806400; 2022YFA0806700), National Natural Science Foundation of China (82430090; 92368105; 82372738; 92157101; 31930022; 32270802), Shanghai Municipal Science and Technology Major Project (2017SHZDZX01, HT), CAS Youth Interdisciplinary Team, and Center for Advanced Interdisciplinary Science and Biomedicine of IHM (QYPY20230034).

## Author contributions Statement

Conceptualization: ZFL, SH, HT, and DW Methodology: ZQL, HRH, LC, WC, CW, YFY, and XY

Investigation: ZQL, HY, HRH, LC, WC, CW, YFY, XY, YW, YMY, CS, YB, HYH, XAW, JC, QC, QW, YX, YC, JC, FL, YL,YD, YS, XFW, JL, XG, and ZT

Visualization: ZQL, HY, HRH, LC, WC, CW, YFY, XY, YW, YMY, and CS,

Funding acquisition: ZFL, SH, HT, RR, CR, DW, YS, and LS Project administration: ZQL, JC

Supervision: ZFL, SH, HT, YH, RR, CR, DW, JL, YS, and LS

Writing – ZQL, HRH, LC, WC, CW, YFY, ZFL, SH, HT, YH, RR, CR, DW, JL, YS, and LS

Writing – review & editing: ZL, SH, HT, YH, RR, CR, DW, JL, YS, and LS

## Figure legend for supplementary figures

**Fig. S1.**
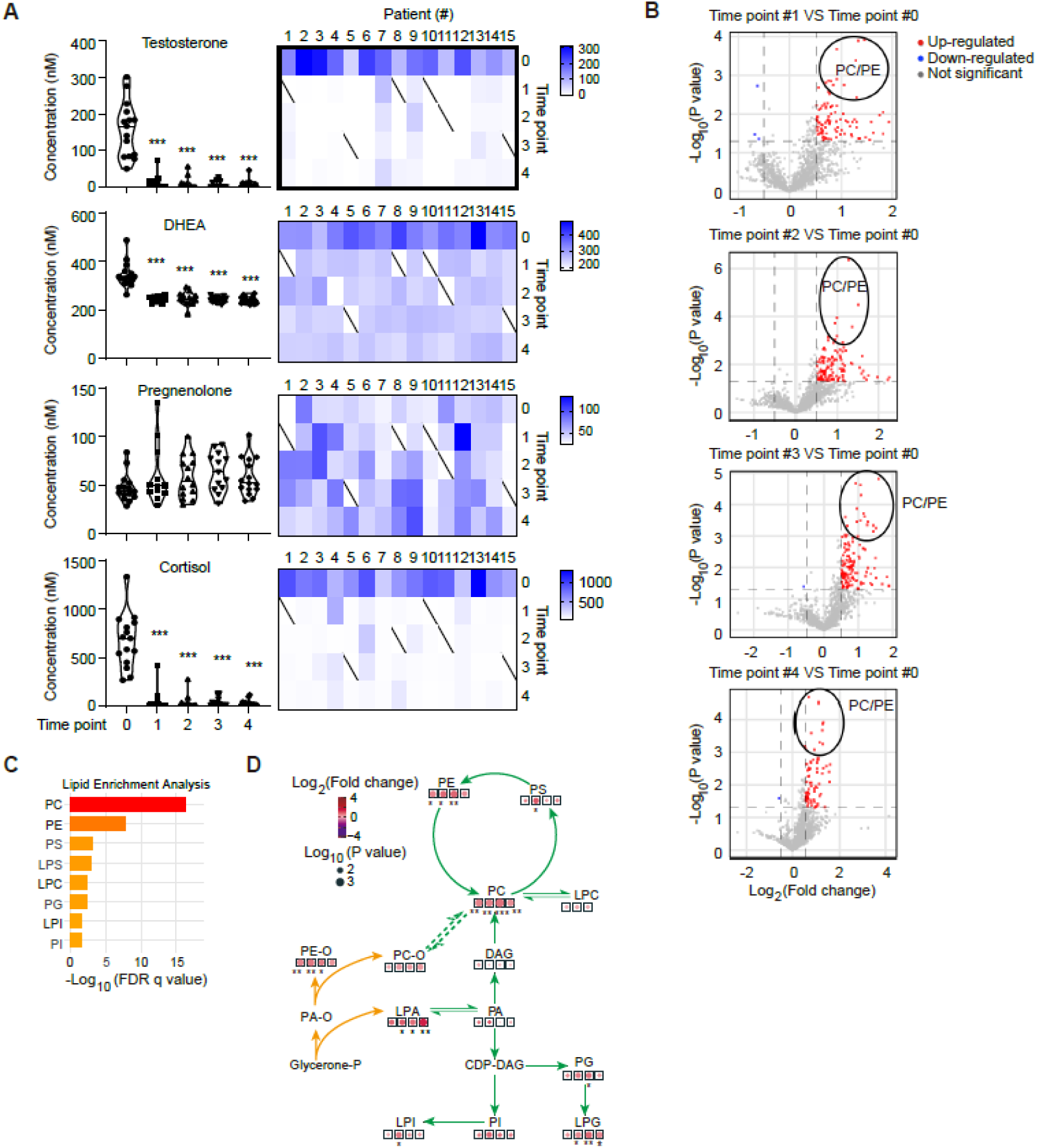
Abiraterone-induced systemic metabolomics alterations in mHSPC patients receiving the combination therapy of ADT and abiraterone plus prednisone (n = 15). **A**, Serum sterol alterations in metastatic hormone-sensitive prostate cancer (mHSPC) patients. For each steroid, the left panel shows aggregate data, and the right panel depicts the individual trajectories for each patient. **B**, Volcano plot of altered lipids in abiraterone-treated mHSPC patients. Significantly upregulated and downregulated lipids are highlighted in red and blue, respectively. **C**, Bar plot of lipid subclasses enriched following abiraterone administration. **D**, A schematic diagram of the phospholipid metabolism pathway. The diagram shows the lipidomic changes across all time points. The four hollow squares (from left to right) represent the comparison of time points #1 through #4 against baseline (time point #0), indicating the direction and consistency of changes in pathway flux. The size of the solid circles corresponds to -log_10_(P value) for the alterations in that particular subclass. The color of the solid circles represents log_2_(Fold change) of lipids in that subclass, providing a visual summary of the impact and statistical confidence. PA, phosphatidic acid; PC, phosphatidylcholine; PE, phosphatidylethanolamine; PG, phosphatidylglycerol; PI, phosphatidylinositol; PS, phosphatidylserine; PA-O, alkylphosphatidic acid; PC-O, alkylphosphatidylcholine; PE-O, alkylphosphatidylethanolamine; LPA, lysophosphatidic acid; LPC, lysophosphatidylcholine; LPG, lysophosphatidylglycerol; LPI, lysophosphatidylinositol; LPS, lysophosphatidylserine; LPE-O, alkyllysophosphatidylethanolamine; CDP-DAG, cytidine diphosphate - diacylglycerol; DAG, diacylglycerol; Glycerone-P, glycerone phosphate; SM, sphingomyelin. *, ** and *** denoted P < 0.05, P < 0.01 and P < 0.001, respectively.

**Fig. S2.**
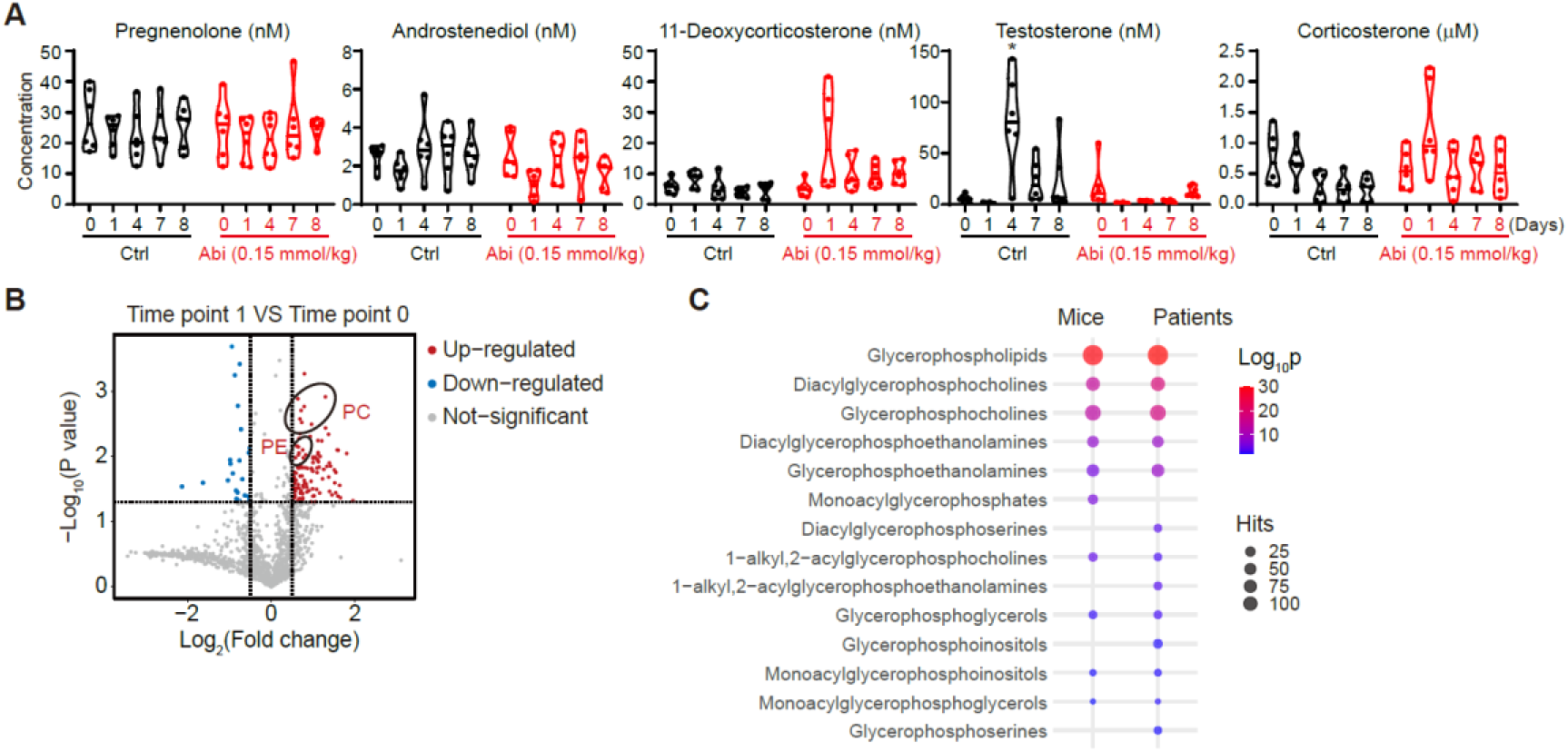
Abiraterone-induced systemic metabolomics alterations in mice. **A**, Violin plots of the serum levels of steroids in abiraterone-treated mice (n = 6 per group). **B**, Volcano plot of altered lipids in abiraterone-treated mice. Significantly upregulated and downregulated lipids are highlighted in red and blue, respectively. **C**, Cross-species conservation of abiraterone-altered lipids in mice and patients. The size of the circles corresponds to the number of significantly changed lipid species within the subclass, and the color represents the log_10_ (P value). *, ** and *** denoted P < 0.05, P < 0.01 and P < 0.001, respectively.

**Fig. S3.**
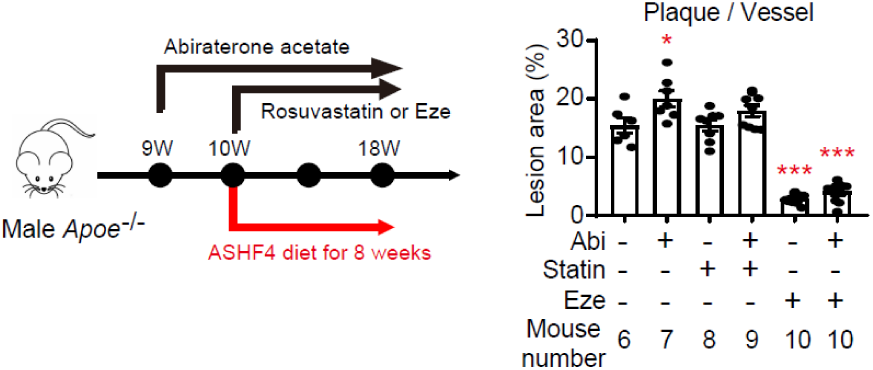
Rosuvastatin and ezetimibe suppress abiraterone-induced atherosclerosis in ASHF4 diet fed male *Apoe* ^-/-^ mice. Ctrl group, n = 6; Abi group, n = 7; Rosuvastatin group, n = 8; Rosuvastatin + Abi group, n = 9; n = 10 for other groups. Eze, ezetimibe, 10 mg/kg/day, per os; Statin, rosuvastatin calcium, 5 mg/kg/day per os. Data are shown as the mean ± SEM. *, ** and *** denoted P < 0.05, P < 0.01 and P < 0.001, respectively.

**Fig. S4.**
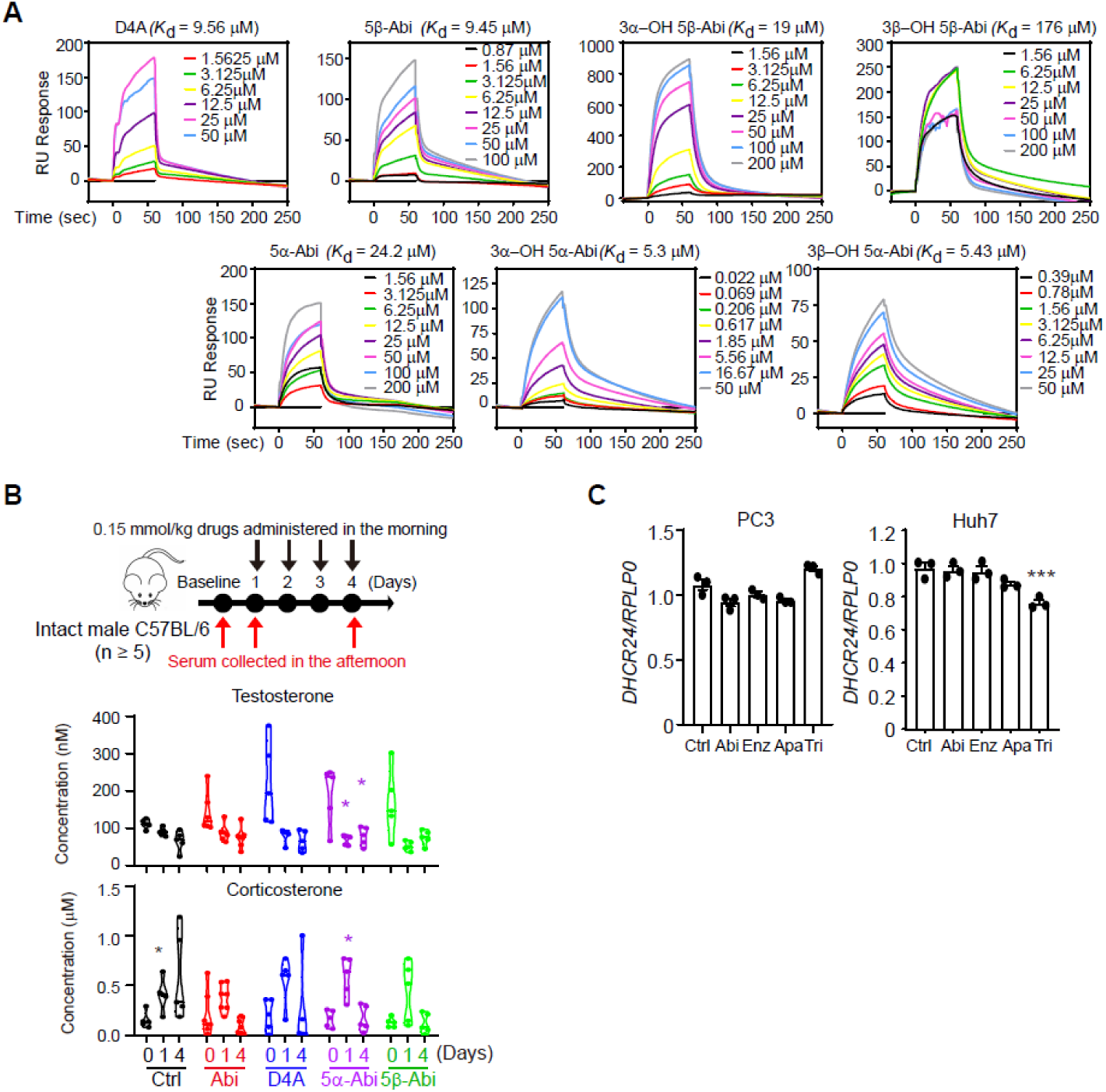
Effect of abiraterone on DHCR24. **A**, Surface plasmon resonance (SPR) analysis on the affinity of abiraterone metabolites to purified DHCR24 proteins. D4A, delta-4 abiraterone; 5α-Abi, 5α-reduced abiraterone; 5β-Abi, 5β-reduced abiraterone. **B**, Intact male C57BL/6 mice were treated with abiraterone acetate or its metabolites (0.15 mmol/kg/day; Intraperitoneal injection). Serum was collected after one or four days post treatment (Abi group, n = 7; n = 5 for other groups). **C**, Abiraterone has limited effect on DHCR24 mRNA abundance. Compounds of 1 µM were used to treat different cell lines for 24 hrs. Data are shown as the mean ± SEM. *, ** and *** denoted P < 0.05, P < 0.01 and P < 0.001, respectively.

**Fig. S5.**
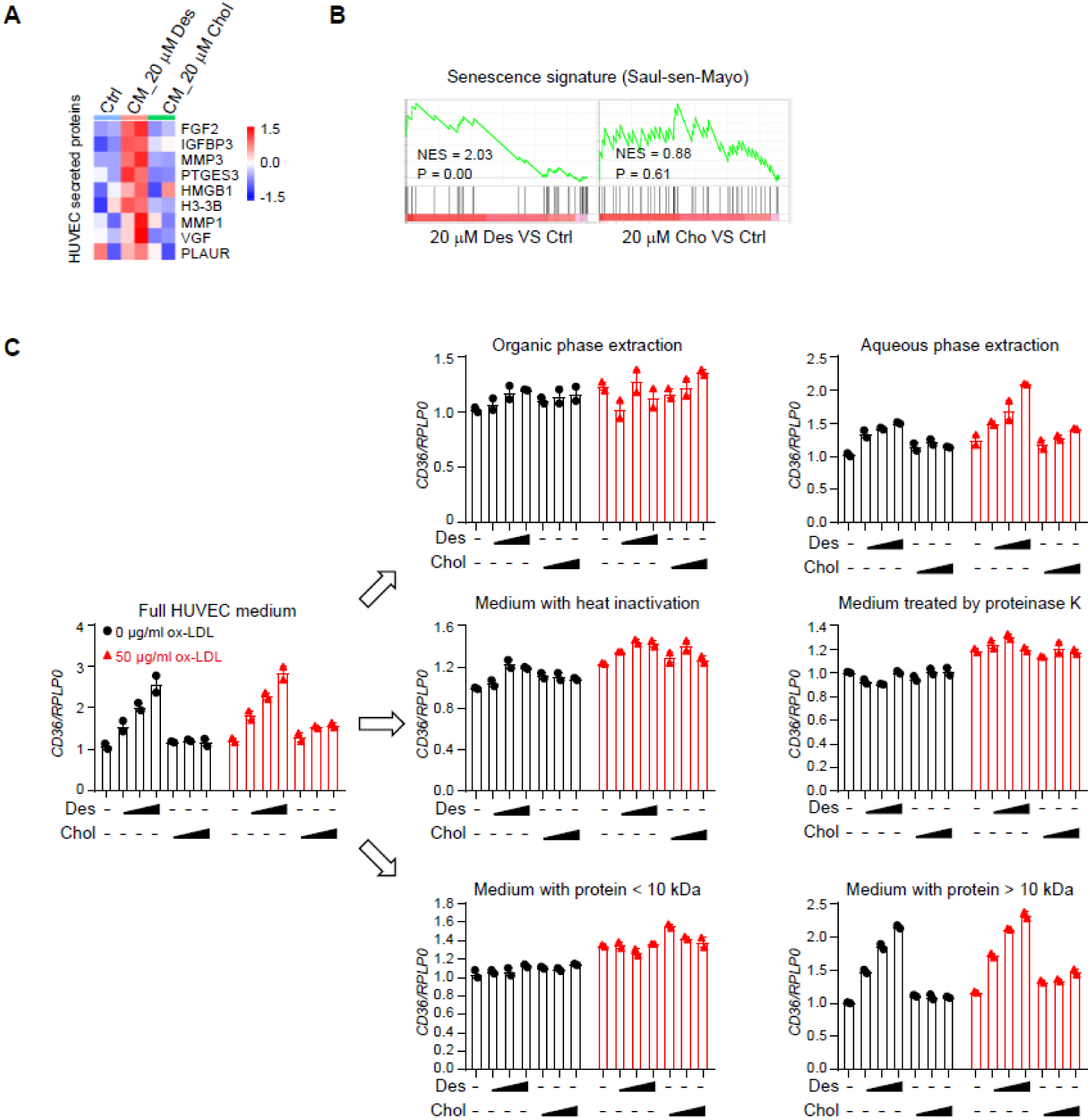
Characterization of a senescence-associated secretome from HUVECs. **A**, Enrichment of senescence-associated secretory phenotype (SASP) related proteins in desmosterol-treated HUVEC CM. HUVECs were treated with 20 μM desmosterol or 20 μM cholesterol for 72 hrs before the medium was collected for proteomics analysis. CM_20 μM Des, conditioned medium from HUVECs treated by 20 μM desmosterol for 72 hrs; CM_20 μM Chol, conditioned medium from HUVECs treated by 20 μM cholesterol for 72 hrs. **B**, GSEA on the proteomic data from HUVEC CM. Saul_sen_Mayo gene set is used as a senescence signature. **C**, Identification of bioactive fraction in desmosterol-treated HUVEC conditioned medium (CM) that upregulates CD36 in THP-1 cells. HUVEC CM was fractionated by MTBE liquid-liquid extraction, heat inactivation (100°C, 10 min)/ protease K digestion (1 mg/mL, 37°C for 2 h), or ultrafiltration (10 kDa MWCO), to determine the activity component in HUVEC CM.

**Fig. S6.**
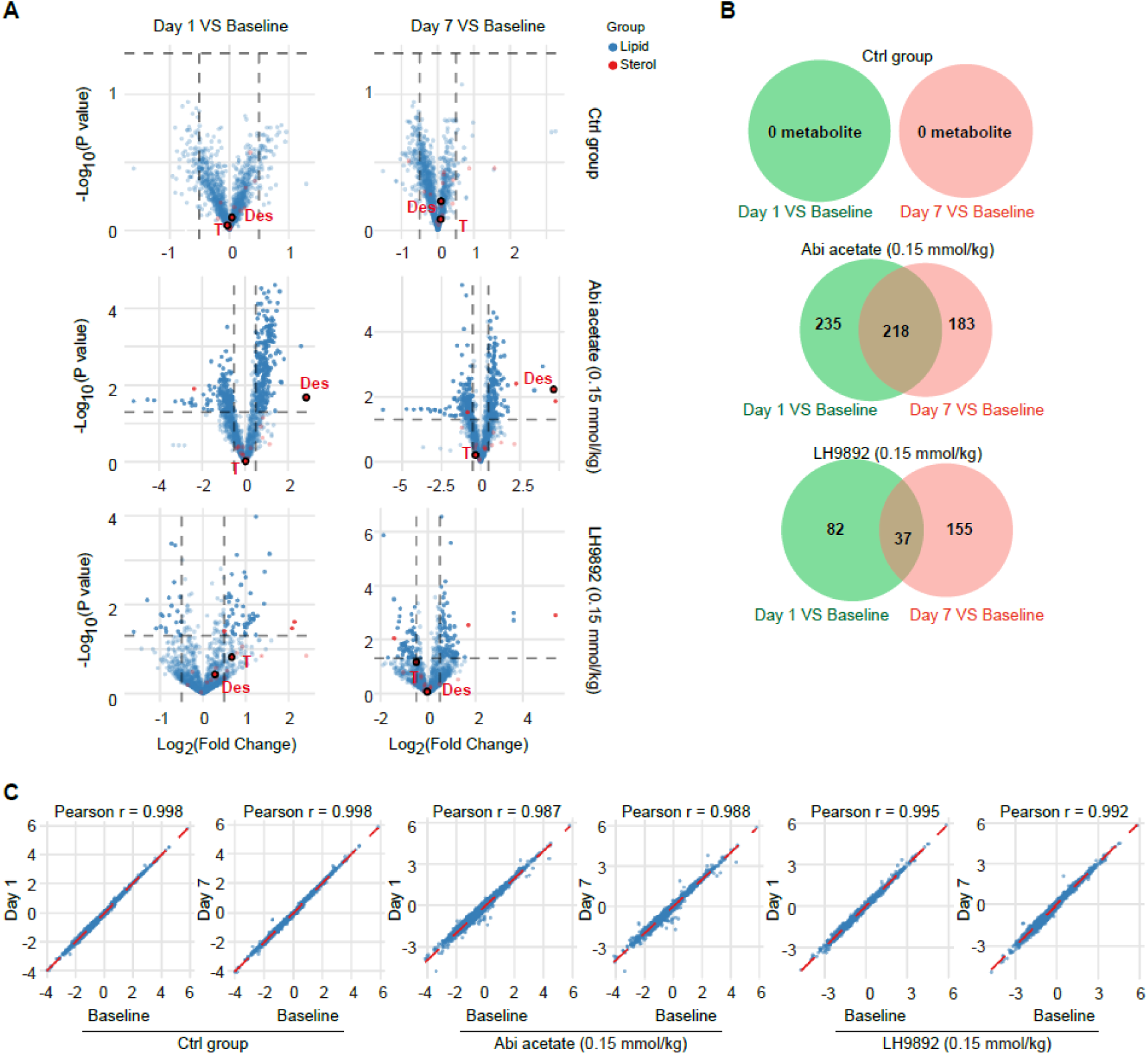
Systemic alterations of lipids and sterols in mice treated with abiraterone acetate or LH9892. **A**, Volcano plot of altered lipids and sterols in control mice (n ≥ 5), abiraterone acetate-treated mice (n ≥ 5), and LH9892-treated mice (n ≥ 5). Intact C57BL/6 mice were treated with abiraterone acetate (0.15 mmol/kg/day, intraperitoneal injection) or LH9892 (0.15 mmol/kg/day, intraperitoneal injection) for 7 consecutive days. Serum samples were collected at the baseline, one day and seven days post treatment. Des, desmosterol; T, testosterone. **B**, Venn diagram showing the numbers of significantly altered metabolites in different mouse groups. **C**, Correlations of metabolites detected at different time points after various treatments.

